# Corticospinal neurons encode complex motor signals that are broadcast to dichotomous striatal circuits

**DOI:** 10.1101/2020.08.31.275180

**Authors:** Anders Nelson, Brenda Abdelmesih, Rui M Costa

## Abstract

Sensorimotor cortex controls movement in part through direct projections to the spinal cord. Here we show that these corticospinal neurons (CSNs) possess axon collaterals that innervate many supraspinal brain regions critical for motor control, most prominently the main input to the basal ganglia, the striatum. Corticospinal neurons that innervate the striatum form more synapses on D1-than D2-striatal projection neurons (SPNs). This biased innervation strategy corresponds to functionally distinct patterns of termination in spinal cord. CSNs are strongly driven during a striatum-dependent sequential forelimb behavior, and often represent high level movement features that are not linearly related to kinematic output. Copies of these activity patterns are relayed in a balanced fashion to both D1 and D2 projection pathways. These results reveal a circuit logic by which motor cortex corticospinal neurons relay both kinematic-related and unrelated signals to distinct striatal and spinal cord pathways, where postsynaptic connectivity ultimately dictates motor specificity.

**Highlights:**

- Corticospinal neurons send axon collaterals most abundantly to the striatum
- Biases in striatal innervation correspond to biases in spinal innervation
- CSNs represent complex movement sequence information
- Corollary motor sequence signals are relayed to both striatal projection pathways

**eTOC Blurb:** Nelson, A. et al. detail the organization of corticospinal neurons and their coordinated cell type-specific targets in the dorsolateral striatum and spinal cord. Corticospinal neurons encode both kinematic-related and unrelated signals during motor sequences, and relay this information in a balanced fashion to dichotomous striatal pathways.

## Introduction

Voluntary movement emerges from neuronal activity distributed across a wide range of motor control structures (Arber and Costa, 2018; Sherrington, 1906). Corticospinal neurons (CSNs), the principle output pathway of sensorimotor cortex, relay command signals to the spinal cord, where their main axons synapse on several distinct classes of spinal interneurons that pattern motor output and shape sensory feedback (Illert et al., 1976, 1977; Lloyd, 1941; Porter and Lemon, 1993; Shinoda et al., 1976; Ueno et al., 2018). Despite their structural primacy, little is known about how CSNs encode motor output, especially during complex behaviors, such as sequences of movements. Ostensibly, CSNs should be active at each of the movements comprising a sequence, a representation that would reflect the conventional perspective that CSNs linearly encode motor output (Porter and Lemon, 1993). Yet, some reports have revealed electrophysiological complexities and plasticity mechanisms of CSNs that suggest their role in controlling movement may be more nuanced (Peters et al., 2017; Ueno et al., 2018). CSNs also give rise to axon collaterals that form synapses in a broad range of brain structures, affording them remarkable – yet largely uncharted – influence over nearly all levels of the motor control neuraxis (Donoghue and Kitai, 1981; Hooks et al., 2018; Kita and Kita, 2012; Ramón y Cajal, 1909). How these corollary synapses in the brain are organized, and what information is transmitted through their activity, remains obscure. Because collaterals of CSNs have the capacity to influence so many brain regions, characterizing the anatomical and functional properties of CSNs is imperative.

In this study, we reveal the wide range of supraspinal brain regions innervated by CSNs, and discover that the striatum is the most innervated of these regions. The striatum is composed of two molecularly distinct populations of spiny projection neurons (SPNs) defined in part by the expression of dopamine receptor type 1 (D1) or type 2 (D2) (Beckstead and Kersey, 1985; Gerfen et al., 1990; Gertler et al., 2008; Graybiel, 1990; Miyachi et al., 1997). While the relative contributions of D1 and D2 SPNs to motor output is a subject of continued study, the coordinated activity of both populations is necessary for many behaviors, including the learned sequencing of body movements, like sequences of lever presses in trained rodents (Cui et al., 2013; Jin and Costa, 2010; Jin et al., 2014; Pisa, 1988). The role of striatum in movement sequences is reflected in the activity of SPNs. For instance, the activity of some SPNs appears to faithfully and directly encode motor output, while other SPNs develop responses not explicitly related to body kinematics, like the onset or offset of lever press sequence rather than individual lever press events (Carelli and West, 1991; Crutcher and Delong, 1984; Jin et al., 2014). Moreover, different fractions of D1 and D2 SPNs display onset and offset responses. What neural structures might contribute to the diversity of SPN sequence encoding properties? One possibility is that motor cortex relays efference copies of motor commands to the striatum, where behavioral state information, sensory information, and reward-related feedback shape SPN activity through intra-striatal and basal ganglia feedback circuits (Houk and Wise, 1995; Redgrave et al., 1999). In this perspective, CSNs are often thought to transmit kinematic information, while intratelencephalic (IT) corticostriatal neurons are thought to transmit higher order task-related information. Indeed, many corticostriatal neurons are active during motor tasks, with pronounced diversity from neuron to neuron (Turner and DeLong, 2000).

Despite these studies, the anatomical and functional properties of CSNs and their projections to striatum remain obscure. Do CSNs only represent kinematic information, while sequence-related activity is limited to IT neurons? Critically, given the fact that D1 and D2 SPNs display different types of sequence-related activity, does the information represented by cortical neurons that synapse on D1 or D2 neurons differ? Addressing these questions has been challenging, partly owing to technical difficulties in monitoring and manipulating cortical neurons defined by their structural and cell type-specific targets in spinal cord and striatum. This functional obscurity is accompanied by an elusive anatomical organization. For instance, D1 and D2 SPNs are completely interspersed in striatum, with little topographical organization or segregation one might leverage using traditional neurotracing methods (Gerfen, 1992). Moreover, while D1 and D2 SPNs receive partially distinct presynaptic input, how CSNs and other cortical subpopulations interface with D1 and D2 SPNs is unclear (Kress et al., 2013; Lei et al., 2004; Wall et al., 2013). Finally, while researchers have to some extent successfully detailed the modularity and developmental specificity of spinal circuits, only recently have genetic and viral tools matured sufficiently to capture and manipulate large populations of corticospinal neurons (Bikoff et al., 2016; Clarke, 1851; Peters et al., 2017; Reardon et al., 2016; Rexed, 1954).

In this study we overcame these technical limitations by first using improved intersectional viral tracing methods to map the brainwide targets of CSNs, revealing striatum as the preeminent target. We then showed using optogenetics-assisted circuit mapping that CSNs innervate both D1 and D2 SPNs, with a bias to the direct pathway. Transsynaptic rabies tracing experiments revealed that CSNs with synapses on D1 or D2 SPNs project to different compartments of cervical spinal cord and synapse on multiple distinct spinal interneuron subtypes. Finally, we used two-photon imaging combined with transsynaptic tracing to show that CSNs can encode information related to both kinematics and behavior sequences, and that this information is transmitted in a balanced fashion to both D1 and D2 striatal pathways.

## Results

### Corticospinal neurons project widely throughout the brain, and most prominently to striatum

Corticospinal neurons possess axon collaterals that form synapses throughout the brain, but the degree to which CSNs innervate each target structure was unclear. We combined intersectional viral expression of fluorescent makers with unbiased anatomical reconstruction in an attempt to quantify the relative innervation of brain regions by axon collaterals of CSNs. First, we labeled cellular inputs to the spinal cord by injecting a retrogradely-transported adeno-associated virus encoding Cre recombinase fused to RFP (AAV-retro-Cre.RFP) into right cervical spinal segments C3-C7, which contain the spinal circuits responsible for forelimb muscle control. In the same animals, we injected a Cre-dependent AAV encoding GFP (AAV-FLEX-GFP) into forelimb-control regions of left sensorimotor cortex, resulting in expression of GFP exclusively in CSNs and their axons throughout the nervous system (Figure 1A-O). We then imaged antibody-enhanced GFP and RFP labeling, and used machine learning based methods to distinguish cell bodies and processes, and map their positions to a common brain atlas (Figure 1P-S). Using this approach, we first noted the widespread and diverse brain regions that project to cervical spinal cord, spanning all levels of the motor neuraxis (Figure S1A-D). Perhaps surprisingly, isocortical structures dominated, comprising 47 percent of the total cellular input (Figure S1C inset, 47±0.03%, N=3). Imaging GFP^+^ labeled (i.e. CSN) axons in the spinal cord revealed widespread varicosities around cervical spinal injection sites, but also substantial collateralization in distant thoracic, and to a lesser degree, lumbar segments (Figure 1B-D). Quantification of GFP^+^ cellular labeling revealed these axons arose from neurons in deep layers of sensorimotor cortex (Figure 1T); this labeling was consistent at the mesoscale across animals (Figure 1T inset, correlation coefficient: 0.98±0.01). CSN axonal labeling in the brain revealed axonal processes in many important forebrain, midbrain, and hindbrain regions, several of which are themselves implicated in motor control (Figures 1T-U, S1F, correlation coefficient: 0.93±0.003). Notably, CSNs project most prominently to the dorsolateral striatum (DLS; Figure 1U, inset, 9.63±0.69% of all neurites), and form abundant synapses in this region as confirmed using synaptophysin-fused fluorescent reporters (Figure S1G-K). Because direct cortical injections of AAV-FLEX-GFP capture GFP-labeled CSNs only around the injection area, we sought to confirm our results using an unbiased intersectional approach to label CSNs that project to DLS (CSNs_DLS_). AAV-retro-Cre.RFP was injected into right cervical spinal cord, and AAV-retro-FLEX-GFP was injected into left DLS, resulting in Cre-mediated recombination in cortical inputs to striatum that also project to spinal cord, regardless of their cortical origin (Figure S2A). Quantifying all axonal projections from these CSNs_DLS_ revealed this population projects throughout the brain, and indeed sends the largest fraction of axons to DLS (Figure S2B-E, N=3). We followed these output mapping experiments by determining the sources of input to CSNs_DLS_. To this end, we used an intersectional transsynaptic approach to drive expression of two viral constructs specifically in CSNs with synapses in striatum: one encoding the avian receptor for EnVA glycoprotein, and the other encoding the rabies glycoprotein necessary for transsynaptic spread (Figure S3). Two weeks later, we injected motor cortex with the pseudotyped, G-deficient rabies construct EnVa-N2cΔG-tdTomato. This construct infects those neurons expressing TVA, and in a subset of those also expressing N2cG, infects and labels synaptic inputs to those neurons (Figure S3B-O) (Reardon et al., 2016). Using anatomical reconstructions, we found isocortical regions like S1 and M2 provide the main source of input to CSNs_DLS_ (Figure S3P, N=3). Surprisingly, the thalamus predominated non-cortical input to CSNs_DLS_, (Fig S3P, inset). These anatomical experiments highlight the capacity for CSNs to influence diverse brain regions involved in motor control, most notably the input nucleus to the basal ganglia.

**Figure 1.**
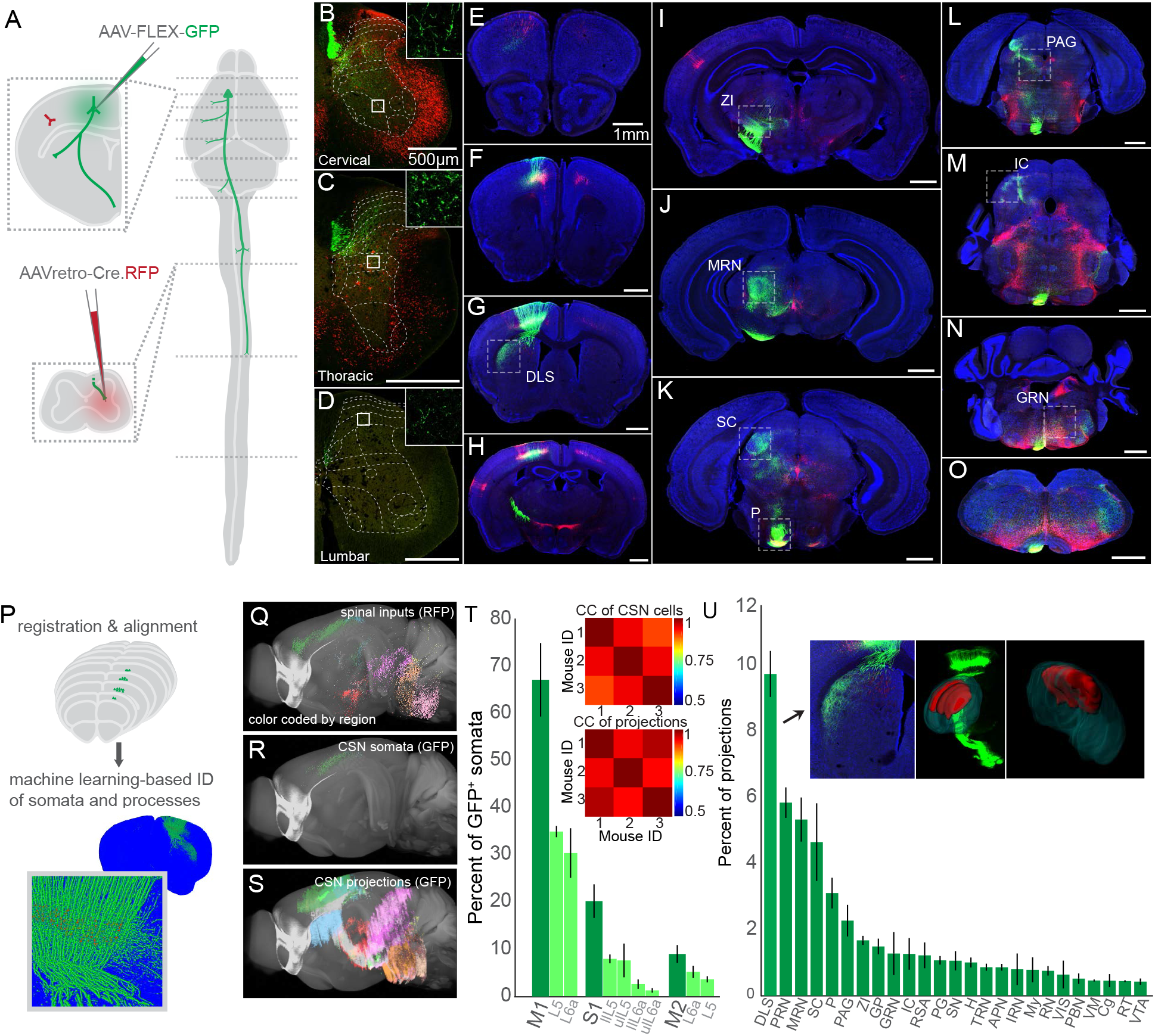
Anatomical characterization of corticospinal neurons. (A) Schematic illustrating the viral injection sites in motor cortex and the spinal cord, and their relative positions in the nervous system. Dashed lines indicate the position of representative images to follow. (B-D) Confocal micrographs of corticospinal neuron (CSN) axons expressing GFP (green) in transverse cross-sections of cervical (B), thoracic (C), and lumbar (D) spinal cord. The insets are high magnification images of GFP^+^ bulbous varicosities from different laminae of cervical (7Sp/8Sp), thoracic (7Sp/ICl), and lumbar (4Sp) segments. Neuronal processes expressing Cre.RFP are in red. (E-O) Confocal micrographs of transverse sections throughout the brain, illustrating GFP^+^ CSNs and their axonal projections (green), along with all spinal inputs made to express Cre.RFP (red). Some regions of interest are boxed by dashed lines and include: dorsolateral striatum (DLS), zona incerta (ZI), midbrain reticular nucleus (MRN), superior colliculus (SC), pons (P), periaqueductal grey (PAG), inferior colliculus (IC), and gigantocellular reticular nucleus (GRN). DAPI is in blue. (P) Illustration of the registration and alignment workflow. A machine learning-based approach was used to identify and mask cell bodies (red) and neurites (green) in registered and aligned coronal images. (Q-S) Three dimensional reconstructions of spinal inputs (Q), CSN somata (R), and CSN neurites (S) throughout the brain. Colors correspond to major brain divisions in which they reside. Note that caudal brainstem is not included in these analyses. (T) The cortical regions giving rise to corticospinal somata. Dark green bars represent the major regions; light green bars represent subdivisions of those cortical regions. The insets show cross-correlation analyses of animal-to-animal CSN somata settlement (above) and CSN processes settlement (below). (U) Top brain regions to which CSNs project, measured as what fraction of all neurites are found within those brain structures, excluding sensorimotor cortex and fiber tracts. The left inset is a high-magnification micrograph of DLS. The middle inset shows Imaris 3D reconstructions of DLS (dark green) and CSN axonal labeling in DLS (red) superimposed over a 3D projection of GFP labeling. The right inset is a caudomedial view of 3D reconstructions from (B). Error bars are SEM.

### CSNs_DLS_ synapse on distinct striatal pathways

Within the striatum, CSNs_DLS_ have the capacity to synapse on two interspersed populations of spiny projection neurons, defined in part by expression of either dopamine receptor 1 or 2 (D1 or D2 SPNs). Previous research revealed that stimulating pyramidal tract-projecting (PT) neurons drives larger currents in D1 SPNs than D2 SPNs (Kress et al., 2013). Yet, PT neurons may also include non-corticospinal populations, including corticobulbar neurons, raising the question of whether CSNs similarly target both D1 and D2 SPNs, and to differing degrees. To address this possibility, we combined an intersectional optogenetic expression strategy with whole-cell voltage clamp recordings to characterize the synapses made by CSNs onto D1 and D2 SPNs. First, we expressed channelrhodopsin-2 (ChR2) in CSNs by injecting AAV-retro-ChR2.tdTomato in cervical spinal cord of adult D1-tdTomato or D2-GFP reporter mice (Figure 2A). Weeks later, we made targeted whole-cell recordings from D1 and D2 SPNs in brain slices, identified in part by the presence or absence of reporter gene expression in cell bodies visualized under DIC optics (Figure 2B-H). Recordings were made from neighboring (within 50um) D1 and D2 SPNs in sequence (N=3, n=12 pairs), or in a subset of experiments, simultaneously (N=3, n=8 pairs). Brief (10ms) photostimulation of ChR2-expressing CSN axons drove excitatory postsynaptic currents (EPSCs) in both D1 and D2 SPNs when measured at membrane holding potentials of −70mV (Figure 2I). Comparing ChR2-evoked currents and charge in pairs of D1 and D2 SPNs revealed that CSN collaterals generate larger responses in D1 SPNs when compared to D2 SPNs (Figure 2J-L, 33.44±5.69 pA for D1, 17.79±3.67 pA for D2, p=0.0037; 7.17±1.12 nC for D1, 3.90±0.871 nC for D2, p=0.006), consistent with what is observed in the broader PT population (Kress et al., 2013). Repeating these experiments using stimulation of intratelencephalic (i.e. non-corticospinal) axon collaterals resulted in equivalently-sized EPSCs in D1 and D2 neurons, suggesting biased innervation of D1 SPNs might be unique to CSNs (Figure S4).

**Figure 2.**
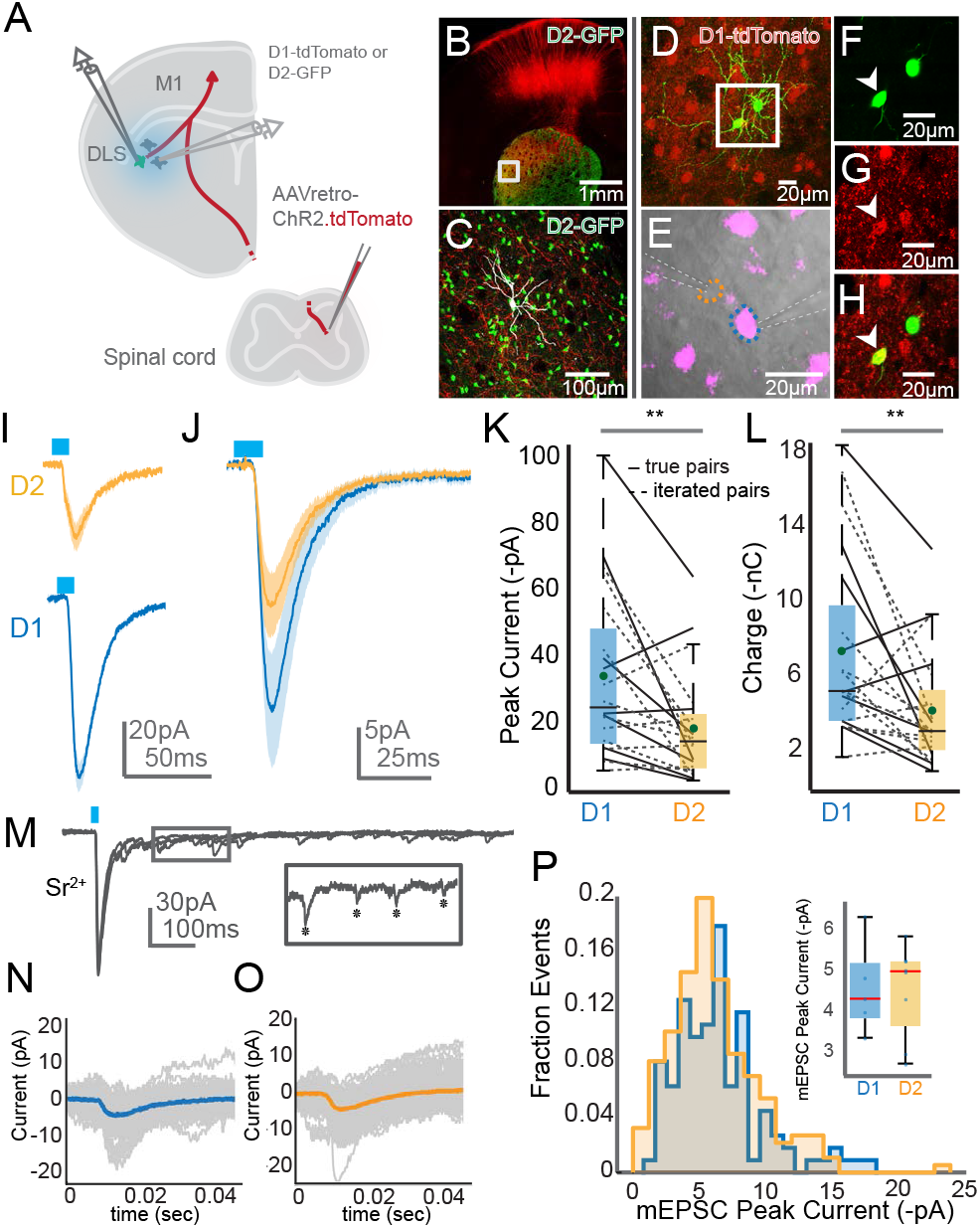
Optogenetics-assisted mapping of corticospinal collateral synapses in the striatum. (A) Schematic of the experimental strategy. (B) Confocal micrograph of a brain slice from an experimental preparation, showing ChR2.tdTomato labeling (red) and transgenically-labeled D2 SPNs (green). (C) High magnification view of the boxed region from (B). An SPN targeted for whole cell recording and filled with neurobiotin is shown in grey. Note the density of CSN axons (red) coursing throughout the recording site. (D) Confocal micrograph of two adjacent SPNs targeted for simultaneous whole cell recordings (green). Transgenically-labeled D1 SPNs are in red. (E) DIC image of one D1 SPN (magenta, blue outline) and one D2 SPN (orange outline) targeted for simultaneous recording. Recording electrode positions are indicated with dashed white lines. (F-H) High magnification single Z plane micrographs from (D) showing GFP (F), tdTomato (G), and overlay (H) fluorescence. The arrowhead indicates a D1 SPN. (I) Whole cell voltage clamp recordings from a D1 SPN (blue) and D2 SPN (orange) in response to optogenetic stimulation (blue bar) of ChR2-expressing axons. Holding potential is −70mV; shaded region indicates SEM. (J) Grand average response of all D1 (blue) and D2 (orange) SPNs to optogenetic stimulation. (K-L) Pairwise comparison of ChR2-evoked amplitude (K) and charge (L) in D1 versus D2 SPNs, recorded either simultaneously (solid lines) or in sequence (dashed lines). (M) Exemplar voltage clamp recordings from an SPN following ChR2 stimulation. Extracellular calcium is replaced with equimolar strontium to desynchronize synaptic release. The inset shows single mEPSCs, indicated with asterisks. (N-O) Trial average of mEPSC evoked from an example D1 (N) and D2 (O) SPN. Individual trials are in grey. (P) Distribution of all mEPSCs ordered by mEPSC peak current, recorded in D1 (blue) or D2 (orange) SPNs. The inset box-and-whisker plot compares average mESPC amplitude in individual D1 versus D2 SPNs.

Results from the above experiments could be explained by CSNs forming either larger synapses onto D1 SPNs than D2 SPNs, or potentially more numerous, but similarly sized synapses. To disambiguate between these possibilities, we replaced extracellular calcium with the divalent cation strontium, which acts to desynchronize neurotransmitter release from the pre-synapse (Figure 2M) (Xu-Friedman and Regehr, 2000). We reasoned that measuring the amplitude of isolated miniature EPSCs evoked by photostimulation would allow us to infer the size of single synapses made by CSN axons on SPNs (Franks et al., 2011). To this end, the averages of mESPCs recorded from D1 or D2 SPNs were indistinguishable, suggesting that CSNs form similarly-sized synapses on both populations (Figure 2N-P, Figure S3I-L; 4.47±0.51 pA for D1, n=5; 4.45±0.40 pA for D2, n=8, p=0.97, N=5). By extension, we tentatively concluded that CSNs, on average, form more synapses on D1 SPNs than on D2 SPNs. Together, these electrophysiological experiments reveal a synaptic and circuit basis by which CSNs interact with two distinct pathways of the basal ganglia.

### CSNs_D1_ and CSNs_D2_ are distinct and terminate in functionally dissimilar spinal compartments

Do identical CSNs synapse on both D1 and D2 SPNs, or could there be partially distinct populations of CSNs biased to innervate one SPN type over the other? How might such distinct populations differently influence motor output through their main descending axons in spinal cord? To address these questions, we turned to an intersectional rabies tracing strategy to map the spinal projections of CSNs_D1_ and CSNs_D2_. Into DLS of D1-Cre or A2a-Cre mice, we injected a cocktail of AAV-FLEX-TVA and AAV-FLEX-N2cG. We later injected EnVa-N2cΔG-tdTomato into the same site, labeling inputs to D1 or D2 SPNs with tdTomato (Figure 3A). Because the input to DLS that projects to spinal cord is motor cortex, we concluded that any axons found in spinal cord arose from CSNs. We then took high resolution confocal images throughout cervical spinal cord, visualizing antibody-enhanced tdTomato labeling, along with co-expression of vGlut1 in order to identify presynaptic boutons (Figure 3B). We first analyzed the distribution of all CSN_DLS_ synapses along multiple segments of cervical spinal cord, noting the expansive terminal fields formed by this population from C3 to C7 (Figure 3C, N=3 for D1-Cre; N=3 for A2a-Cre). Interestingly, CSN_DLS_ synapses were found all across the dorsoventral aspect of the spinal grey, with densest innervation confined to intermediate and superficial spinal laminae.

**Figure 3.**
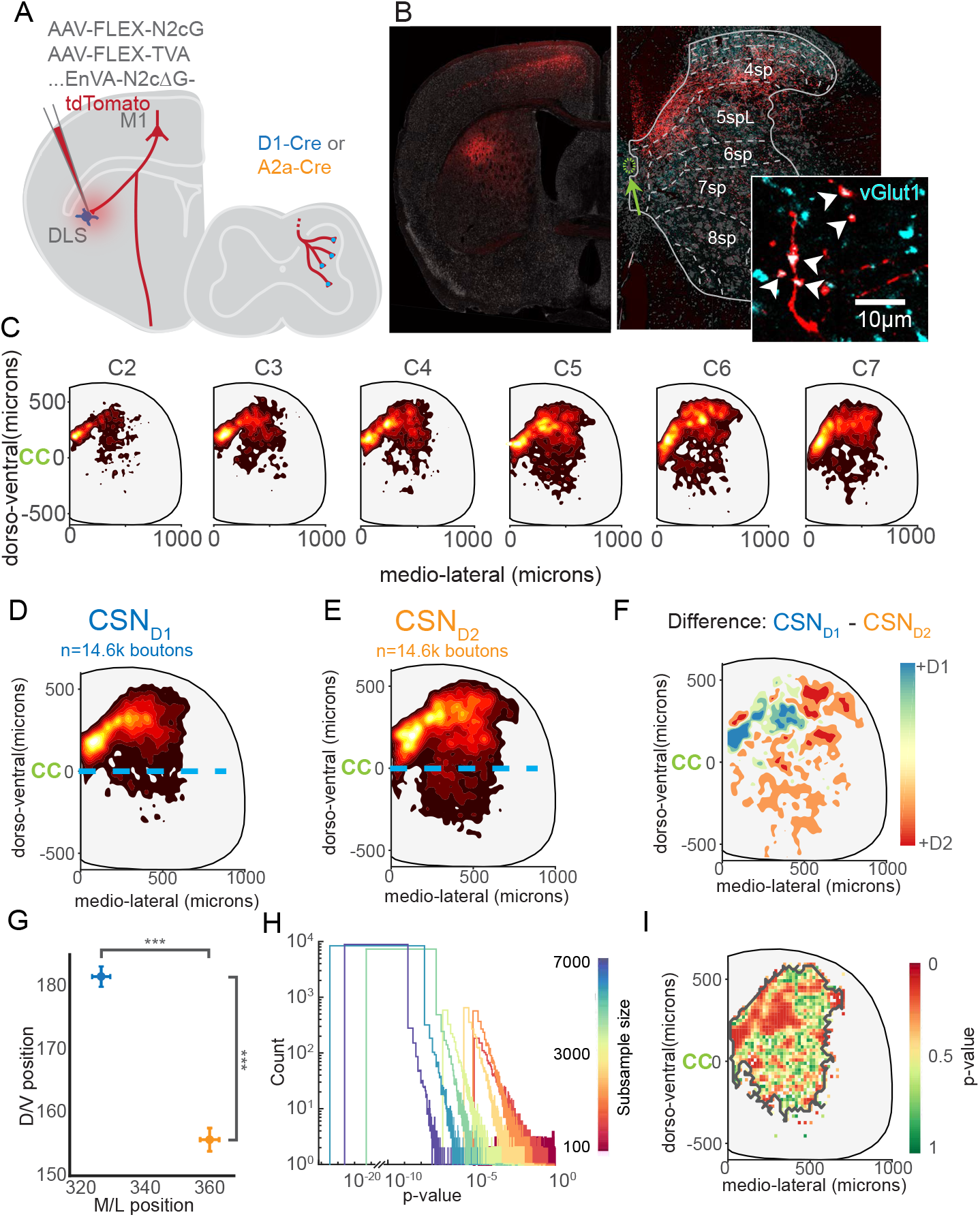
Mapping the distribution spinal synapses from CSNs_DLS_. (A) Experimental strategy to transynaptically label CSNs_D1_ and CSNs_D2_. (B) Photomicrograph of tdTomato-labeled CSNs with identified synapses on striatal SPNs (above), and the synapses formed by these neurons in spinal cord (below). tdTomato^+^ synapses are identified by coincident expression of vGlut (cyan). The green arrow indicates the central canal. Fluorescent Nissl stain is grey. (C) Contour plots illustrating the relative distribution of synapses arising from all CSNs_DLS_, ordered by cervical spinal segment. (D-E) Contour plots illustrating the relative distribution of synapses arising from CSNs_D1_ (D) and CSNs_D2_ (E). (F) The difference between contour plots in (D) and (E). (G) Quantification of the mean mediolateral and dorsoventral settlement of CSNs_D1_ (blue) and CSNs_D2_ (orange). (H) Random resampling analysis. The dataset was resampled with different sample sizes (color), and statistical analysis was repeated many times. The X axis is broken to indicate the heavily skewed distribution. (I) Statistical differences between the spatial distribution of CSNs_D1_ and CSNs_D2_ synapses. See Supplemental Material for details.

We next separately analyzed the distribution of synapses arising from CSNs_D1_ and CSNs_D2_. We found that while both populations of neurons formed synapses spread throughout cervical spinal cord, CSN_D1_ synapses confined to more rostral and medial coordinates (Figure 3D-F). Notably, CSN_D2_ synapses were skewed to the ventral regions of spinal cord, where there is a pronounced settlement of interneuron populations that shape motor output (Figure 3G, 155.75±1.83μm for D1 versus 180.47±1.61 μm for D1, p=3.91×10^−24^)(Bikoff et al., 2016; Briscoe et al., 2000; Eccles et al., 1961). Random subsampling and repeated statistical testing revealed these results were statistically robust, even when sampling less than 6.25% of the total dataset (Figure 3H, i.e. 1100 out of 17,599 resampled coordinates, median of repeated t-tests, dorsoventral: p=0.0065; mediolateral: p=3.5×10^−4^). Finally, we binned coordinates of CSN_D1_ and CSN_D2_ synapses, and performed regional comparative statistics to identify spinal compartments with significantly different innervation patterns. This analysis revealed a large swath of intermediate and superficial laminae with statistically significant innervation differences, as well as smaller hotspots in lateral and ventral regions of the spinal cord (Figure 3I, Figure S5).

These results suggest CSNs_DLS_ have functional access to spatially confined spinal interneurons with distinct roles in motor control and sensory processing. To test this possibility, we performed a series of intersectional transsynaptic rabies tracing to determine if CSNs_DLS_ make synapses on multiple genetically-defined interneuron subtypes. We chose distinct interneuron populations derived from clades defined by expression of unique genetic markers. Specifically, the GABAergic neurons responsible for presynaptic inhibition of proprioceptive feedback are derived from the dorsal class DI4, and express the genetic marker GAD2 (Fink et al., 2014). V2a propriospinal excitatory neurons, which relay copies of forelimb motor commands to the central nervous system, express Chx10 (Azim et al., 2014). Finally, somatostatin-expressing (SST) neurons in dorsal laminae modulate mechanoreceptive sensory feedback (Duan et al., 2014). We first expressed TVA and N2cG in these spinal neurons by injecting AAV-FLEX-TVA and AAV-FLEX-N2cG into cervical spinal cord of GAD2-Cre, Chx10-Cre, or SST-Cre mice (Figure S6). We then injected AAV-FRT-GFP into forelimb motor cortex. Weeks later, we injected EnVA-N2cΔG-FlpO.mCherry into spinal cord, resulting in expression of Flp.mCherry in neurons that synapse on the spinal interneuron of interest, in turn driving expression of GFP in those presynaptic neurons in motor cortex. We confirmed the expression of FlpO throughout the brain by imaging mCherry-labeled neurons, which spanned multiple brain regions, including sensorimotor cortex (Figure S6B-C). A large subpopulation of mCherry-expressing cells in motor cortex co-expressed GFP, indicating FlpO expression successfully drove recombination in CSNs with synapses on spinal interneurons of interest. Consistent with existing results (data not shown), we observed non-uniform distribution of CSN somata across cortical region, depending on target spinal cell type (Figure S6B). Finally, we mapped the position of GFP-labeled neuronal processes throughout the brain using methods described for Figure 1. CSNs with synapses on all interneuron subtypes of interest formed widespread axonal arborizations in DLS (Figure S6D, N=2 for each genotype). While DLS was the recipient of the largest proportion of innervation, we observed significant differences in the proportion of neurites across several brain regions. These results indicate that CSN_DLS_ neurons form synapses on spinal interneuron populations with highly divergent roles in sensory processing and motor control in the spinal cord, and that these subpopulations may exert unique control over supraspinal motor control structures.

### CSNs encode sequence-related activity in a striatal-dependent forelimb task

Compared to other cell types and other brain regions, the activity properties of CSNs in relation to complex behaviors are poorly characterized. The anatomical complexities of CSNs – particularly their prominent projections to the striatum – inspired us to characterize their activity during a behavioral task relevant to basal ganglia: a sequential lever press task. Here, water-restricted mice depress a small lever positioned in front of their right forepaw quickly four times in succession to receive a water reward (Figure 4A). Mice learn this task within several days, as evidenced by the rapid execution of grouped lever presses, and the increased performance of four press sequences and decrease of two press sequences (Figure 4B-D, post hoc t-test, p=0.0284 and p=0.026, respectively, N=8). To analyze kinematic performance with high resolution, we implanted wire electrodes made for recording electromyographic (EMG) signals into four forelimb muscles comprising two antagonist pairs: biceps and triceps, as well as extensor digitorum communis (EDC) and palmaris longus (PL) (Akay et al., 2006). To monitor the activity of CSNs during behavior, we injected retrogradely transported virus encoding GCaMP6f (AAV-retro-GCaMP6f) into right cervical spinal cord of D1-Cre or A2a-Cre mice (Figure 4E-G), and implanted a cranial window over left forelimb motor cortex. Two-photon (2p) imaging was used to record GCaMP6f activity in dendritic trunks of CSNs approximately 300μm below the pial surface (Figure 4H). These dendritic signals are highly correlated with somatic calcium activity, and afford higher temporal resolution than signals from cell bodies (Beaulieu-Laroche et al., 2019; Mittmann et al., 2011; Peters et al., 2017). Calcium signals were extracted using CNMF and highly correlated (rho > 0.8) processes were treated as belonging to the same neuron to minimize overrepresentation by branching dendritic processes (Figure 4I) (Beaulieu-Laroche et al., 2019; Peters et al., 2017; Pnevmatikakis et al., 2016). Aligning Z-scored calcium activity of all neurons from an exemplar mouse to behavior revealed most neurons were strongly active before and during individual lever presses, but with substantial variability from neuron to neuron (Figure 4J-K). To overcome the temporal limitations of analyzing calcium transients, we deconvolved our calcium signals to estimate CSN spiking activity, which we again aligned to lever press sequences. Z scored spiking activity from one mouse (Figure 4L) and across all mice (Figure 4M) was substantially faster than calcium activity, and there was a strong trend for neuronal activity to be enhanced around lever press (Figure 4N). Does peak activity occur at the same time relative to lever press for all neurons? Heatmaps of Z scored activity aligned to single lever press revealed a temporal distribution of peak responses (Figure S7A). Binning neurons by the time of their peak responses revealed most neurons were active immediately after lever press (Figure S7B, median time to peak response ~93ms), with some heterogeneity. Interestingly, the average activity of neurons with peak activity closer in time to lever press was larger than the average activity of neurons with peak responses before or after lever press (Figure 4O).

**Figure 4.**
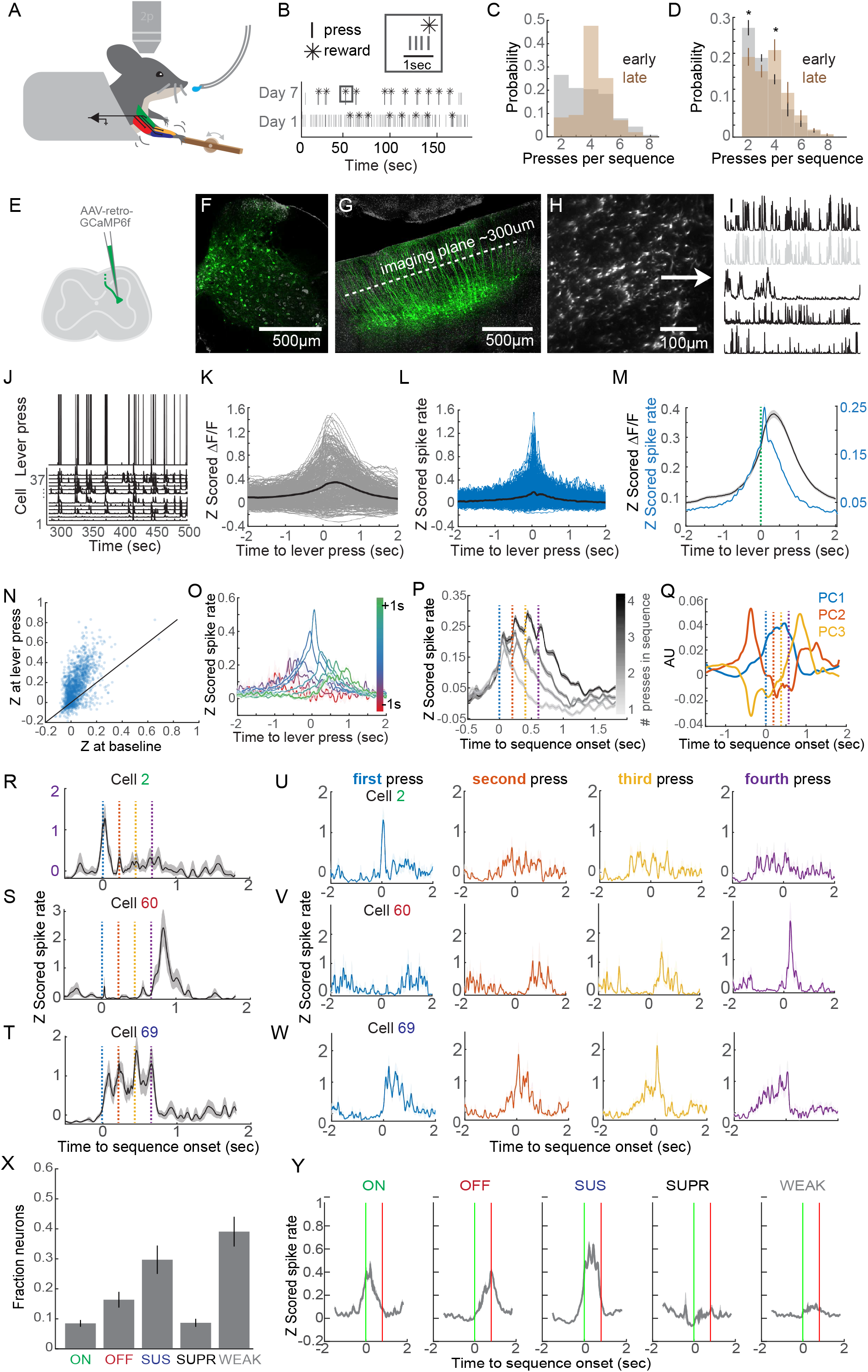
Two-photon calcium imaging of corticospinal neurons during a sequential forelimb behavior. (A) Cartoon of the lever press behavior, two-photon imaging, and EMG recording. (B) Example of lever press sequencing at training day 1 and day 7. Note the development of grouped lever presses in the inset. (C) Probability of sequences containing different numbers of lever presses early in training (grey) and late in training (brown) from a single mouse. (D) Same as C, for all mice. (E) Strategy to label CSNs with GCaMP6f. (F) Confocal micrograph of the injection site. (G) Confocal micrograph of GCaMP6f-labeled CSNs, with the approximate imaging plane indicated. DAPI is in grey. (H) Two-photon image of GCaMP6f expression in cross sections of dendritic processes belonging to CSNS. (I) Example of calcium events derived using CNMF. The second trace is greyed out to indicate it is highly correlated to the top trace, and likely originates from the same cell. (J) Example Z scored calcium activity aligned to lever press events. (K) Trial-averaged calcium activity aligned to lever press for neurons from a single mouse. (L) Same data as (K), but for inferred spiking activity. (M) Average Z scored calcium (grey) and inferred spiking (blue) activity for all neurons and all mice aligned to single lever presses. (N) Z scored spiking activity at rest versus at lever press for all neurons. (O) Average activity traces for neurons with peak activity that falls within different bins of time relative to lever press. (P) Average Z scored spiking activity aligned to the onset of lever press sequences, segregated by sequence length. Inter-press intervals are standardized using time warping, and the press times are indicated with colored dashed lines (Q) The top three PCs of time-warped neuronal activity. (R-T) Examples of neurons with activity coincident with the onset (R), offset (S), or individual presses (T) of lever press sequences. (U-W) The same neurons as (R-T), instead displaying spiking activity aligned to first, second, third, or forth press in the sequence. (X) The fraction of neurons classified as with onset (ON), offset (OFF), sustained (SUS), suppressed (SUPR), or weak (WEAK) activity profiles, across all mice. (Y) The average activity of neurons belonging to each activity profile, aligned to four lever press sequence onset.

How do CSNs encode motor sequences? Does CSN activity linearly relate to motor output by being active around each lever press in a lever press sequence? To answer these questions, we identified and grouped sequences of lever presses of one, two, three, or four presses. To account for variability in the behavior, we used time warping to standardize the inter-press interval within sequences to 200ms, allowing us to preserve temporal resolution when averaging across trials (Figure S8A-B). Aligning activity from the total population of neurons (n=2,374, N=8) to lever press sequences revealed that, on average, CSN activity scales in duration to lever press sequences of increasing length (Figure 4P, Figure S8C-D). This was reflected in the projection of the top three principle components (PCs) of the normalized neuronal activity, which when plotted against each other revealed prominent peaks as neural activity evolved throughout sequence execution (Figure S8E). Yet, when plotted individually, the top three PCs of CSN activity each displayed unique activity signatures. The component accounting for the most variance was elevated in activity throughout sequence execution, while the next two PCs were active most strongly at the onset or offset of sequence. This striking result motivated us to characterize the activity patterns of single neurons. For instance, does the activity of single neurons mirror the population, or is there heterogeneity across cells? To address this possibility, we aligned the time warped Z-scored spiking activity of single neurons to four-lever press sequences. In this way, we were able to visualize the degree to which single neurons are active at different presses in the sequence. Remarkably, we found a heterogenous population of neurons, including those with apparent preferential or selective activity around the first or final press in a sequence, as well as neurons that were robustly active around each press in a sequence (Figure 4R-T), all of which were intermingled in the same fields of view. We complemented this analysis by aligning Z scored activity to the first, second, third, or fourth lever press within a four-lever press sequence, revealing strikingly selective response properties in the same neurons (Figure 4U-W).

Motivated by these results, we next sought to quantify and catalogue the different response properties of individual CSNs. To this end, we aligned binned Z-scored spike rates to time warped lever press sequences, and identified neurons with significant modulation at different windows of the lever press sequence. Using this approach, we identified 8.26% and 16.04% of CSNs responding at the onset (ON) and offset (OFF) of sequence, respectively, as well as a large population of neurons with activity sustained (SUS) throughout sequence execution (Figure 4X, 29.18%). We further identified a population of neurons with activity significantly suppressed (SUPR) relative to baseline (8.45%), as well as population of neurons that did not meet criteria for significant modulation (38.41%). Averaging the Z scored activity of categorized neurons across revealed the relative stereotypy of these responses (Figure 4Y), and this data was recapitulated by inspecting the timing of peak responses of categorized neurons (Figure S9).

### CSN activity is diversely related to muscle activity

Muscle activity may change from lever press to press, raising the possibility that the variability we observe in neuronal activity could be due to differential recruitment of musculature at the onset or offset of sequences. By extension, neuron to neuron response variability may be partially explained by a preferential correlation with individual muscles. To directly address these possibilities, we analyzed the EMG activity of biceps and triceps during behavior and in relation to neuronal activity (Figure 5, Figure S10A-B). First, aligning all biceps activity to local peaks in triceps activity revealed a robust alternation of the activity between these two antagonist muscles (Figure 5A). Biceps and triceps activity alternated immediately preceding lever press, with triceps activity following biceps activity, consistent with the flexor and extensor identity of these muscle groups (Figure 5B). Most importantly, biceps and triceps activity were strongly alternating during lever press sequences, and the amplitude of EMG activity preceding lever press events was similar throughout the sequence (Figure 5C-D). We leveraged our EMG dataset by correlating spike rate of CSNs to biceps and triceps activity during concatenated periods of behavioral quiescence or concatenated lever press sequences. On average, CSNs were more correlated with triceps activity than biceps during random periods of activity and quiescence, but this preference was lost when correlating neural activity with only concatenated lever press sequences, even when controlling for the number of samples in each condition (Figure S10C). We next measured the correlation between average time warped biceps and triceps EMG activity and average time warped spike rate of ON, OFF, SUS, and SUPR neurons during lever press sequences. As expected, SUS neurons were the most correlated with muscle activity, while ON and OFF CSNs were both less correlated with both biceps and triceps activity (Figure 5E). Also as expected, the negative correlation coefficients of SUPR neurons revealed this population was anticorrelated with muscle output. Importantly, on average, no group of neurons was consistently more correlated with biceps or triceps EMG (Figure 5E), suggesting encoding of muscle identity cannot explain sequence encoding properties of CSNs. However, we were able to find many individual neurons strongly correlated with one muscle over the other (Figure 5F). Within this group, neurons with biceps-biased correlation coefficients were on average less strongly modulated during lever press sequence than the population average (Figure S10D-E). Conversely, CSNs biased toward triceps activity were more strongly driven during lever press sequence compared to the population average.

**Figure 5.**
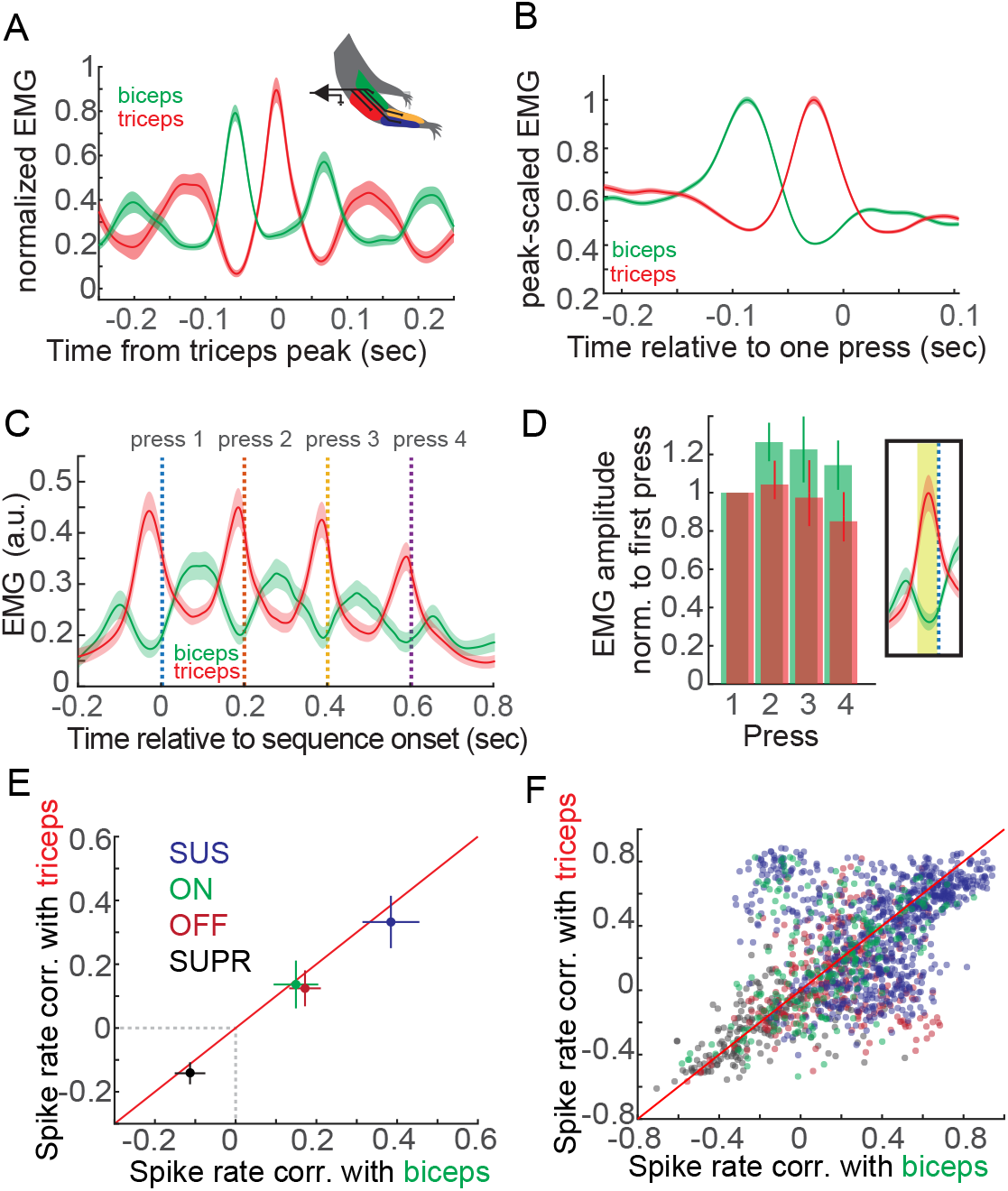
Muscular control of sequential forelimb movements. (A) Example recording of biceps and triceps muscle activity from one mouse. Biceps EMG is aligned to peaks in triceps EMG. (B) Biceps and triceps EMG aligned to single lever press onset, for all mice. (C) Time warped biceps and triceps EMG aligned to lever press sequences, for all mice. (D) Quantification of the mean biceps and triceps activity immediately preceding first, second, third, or forth lever presses in a sequence. The inset indicates the time window for averaging activity. (E) Mean correlation of activity with biceps versus triceps EMG for neurons sorted into ON, OFF, SUS, or SUPR categories. (F) Same as (E), but showing the individual neuron correlations with biceps versus triceps.

### Similarly diverse CSN activity is relayed to both striatal projection pathways

Like CSNs, striatal SPNs can selectively encode the onset and offset of lever press sequences, as well as display sustained activity throughout sequence (Jin and Costa, 2010). We wondered 1) if CSNs with identified synapses in the striatum (i.e. CSNs_DLS_) show similar encoding of lever press sequences, and 2) if this extends to CSNs_DLS_ that innervate D1 or D2 SPNs (i.e. CSNs_D1_ or CSNs_D2_), and if the fraction of classified neurons is similar between these groups. To tackle this challenge, we combined our 2p calcium imaging experiments with *in vivo* transsynaptic rabies tracing from D1 or D2 SPNs. In the same mice as above (i.e. D1-Cre or A2a-Cre), we injected AAV-FLEX-N2cG and AAV-FLEX-TVA into DLS before cranial window implantation (Figure 6A). After all functional calcium imaging data was acquired from these mice, EnVA-N2cΔG-tdTomato was injected into the same location of DLS, using an angled pipette approach, entering the brain caudal to the imaging coverslip (Figure 6B-D). Ten days following rabies injection, we took structural images of tdTomato and GCaMP labeling in motor cortex, as well as Z stacks of tdTomato labeling (Figure 6E, Figure S11A). We then used 3D reconstruction to improve detection of tdTomato^+^ dendrites at the functional imaging plane, and generated binary masks from this dataset. Finally, we used the GCaMP structural reference images to align binary masks of rabies labeling to the functional imaging dataset (Figure S11B). This approach allowed us to identify CSNs_D1_ and CSNs_D2_ post hoc, avoiding any effect rabies expression has on electrophysiological response properties. We first analyzed neuronal activity in CSNs with confirmed synapses in the striatum (i.e. neurons that synapse on either D1 SPNs or D2 SPNs. Grouping this data, we confirmed that the CSN_DLS_ population shows activity that scales in duration with lever press sequences, similar to general CSNs (Figure 6F). Do CSNs_D1_ and CSNs_D2_ comprise similar proportions of ON, OFF, SUS, and NEG neurons as does broader CSN population, and are the proportions of these categories different between inputs to D1 versus D2 SPNs? We applied our classification scheme to rabies-labelled CSNs, and compared these data to unlabeled neurons from the same mice, in an attempt to control for any differences in rabies expression across animals. We found similar proportions of ON, OFF, SUS, and NEG neurons in tdTomato^+^ neurons compared to tdTomato^−^ CSNs, along with no clear enrichment of any classification of neuron type when comparing CSNs_D1_ and CSNs_D2_ (Figure 6G). These results indicate that information encoded by CSNs is transmitted in a balanced fashion to both D1 and D2 SPNs when measured using an anatomical assay.

**Figure 6.**
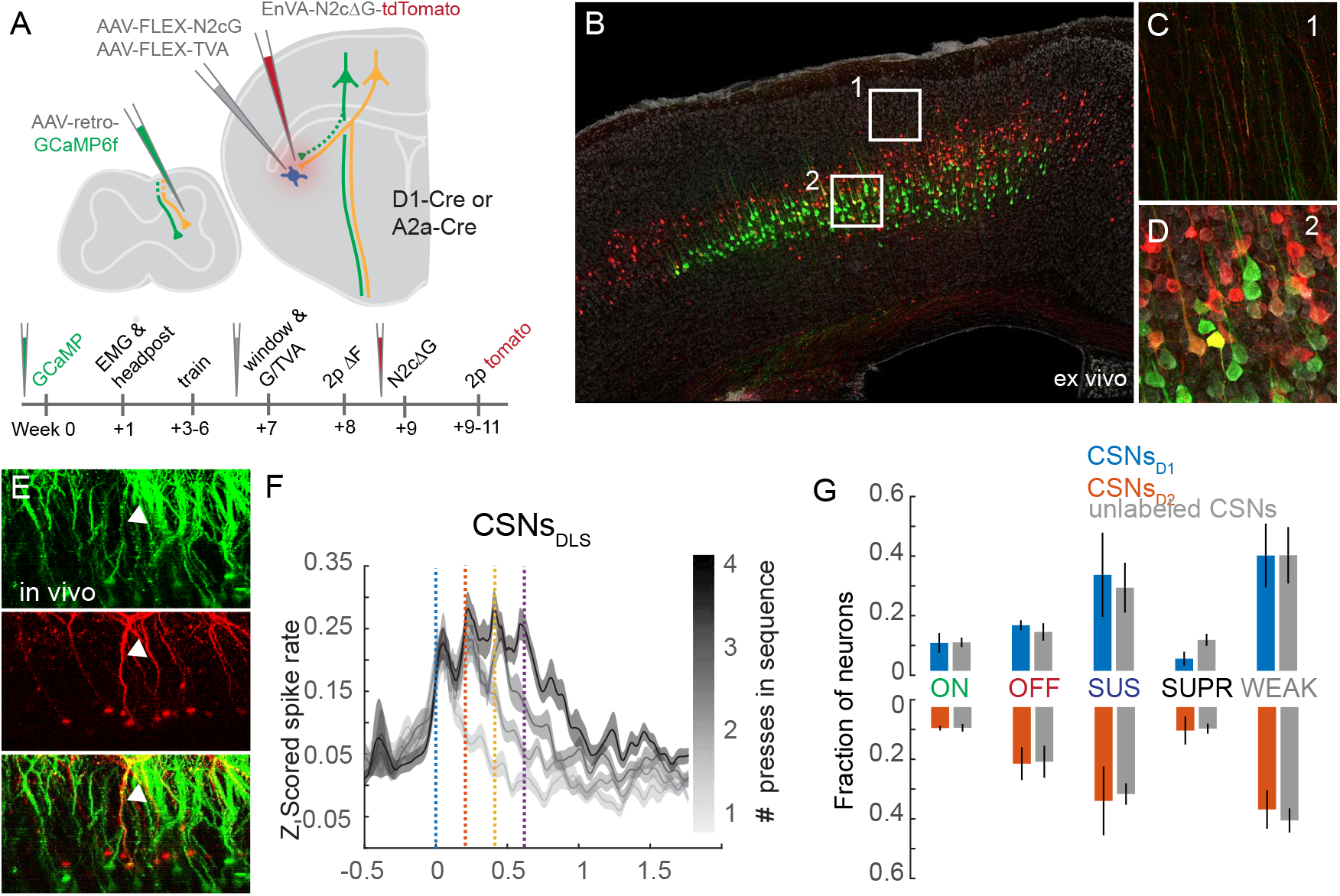
Rabies-based in vivo identification of corticospinal neurons with striatal synapses. (A) The experimental strategy and timeline. (B) Confocal micrograph of GCaMP6f-tagged CSNs (green) and transynaptically-identified inputs to striatal SPNs (red). DAPI is grey. (C-D) High magnification images of the regions indicated in (B) showing green and red fluorescence in dendritic trunks (C) and somata (D). Double-labeled processes are yellow. (E) X-Z view of GCaMP- and tdTomato-expressing neurons in motor cortex, imaged *in vivo*. A double-labeled neuron is indicated with the arrowhead. (F) Average Z scored spiking activity of CSNs_DLS_ aligned to the onset of lever press sequences, segregated by sequence length. (G) Fraction of ON, OFF, SUS, SUPR, and WEAK transynaptically-identified CSNs_D1_ and CSNs_D2_, compared to unlabeled CSNs.

## Discussion

The results presented here reveal that corticospinal neurons encode more than kinematic information, including sequence-related information in the form of onset or offset responses. These results further uncover the structural and functional principles by which this complex corticospinal neuronal activity is transmitted in a balanced manner to spinal and basal ganglia circuits critical for motor control. We used a combination of anatomical and electrophysiological tools to show that CSNs form axon collaterals throughout the brain, but most abundantly in the dorsal striatum, where they form more synapses on D1 SPNs compared to D2 SPNs. This synaptic bias is accompanied by an anatomical divergence: CSNs that synapse on either D1 or D2 SPNs form distinct terminal fields in cervical spinal cord, underscoring their capacity to differentially regulate spinal circuits by synapsing on spatially segregated interneuron populations. Using functional imaging during a skilled motor sequence behavior, we showed that the activity of many CSNs is closely related to muscle activity. Remarkably, a substantial proportion of CSNs showed activity that was not well-explained by muscle output, but instead was tightly correlated with other features of motor sequences. Combining 2p imaging with transsynaptic tracing revealed that these diverse activity profiles are equally represented in CSNs that form synapses on D1 or D2 SPNs. The biased distribution of CSN synapses between D1 and D2 SPNs, as well as their cognate projection patterns in spinal cord, promote a mechanism by which movement-related information is simultaneously broadcast to multiple motor control structures, where differences in postsynaptic connectivity shape neural activity to ultimately direct motor specificity.

### Implications for motor cortical control of movement

Evidence for importance of motor cortex in directing skilled motor output across the animal kingdom is strong. Perturbations to motor cortex abolish or degrade skilled forelimb behaviors, and well-established electrophysiological and analytical methods have revealed computational principles underlying the apparent transformation of cortical activity to command signals resembling muscle output (Fetz, 1993; Guo et al., 2015; Miri et al., 2017; Russo et al., 2018). While such studies position motor cortex as a principle controller of movement, important biological constraints should be considered. First, many efforts to study motor cortex do so irrespective of cellular identity. While array recordings from deep layers of motor cortex probably include a fraction of corticospinal neurons, many more of those unidentified units likely do not project to the spinal cord, and perhaps subserve functions not explicitly related to muscle output. Our calcium imaging approach overcomes this limitation by allowing us to record exclusively from corticospinal neurons, including those with sparse activity that may otherwise be lost with traditional electrophysiological methods. Next, in adult rodents and other animals, skeletal motor neurons receive only indirect input from motor cortex, mediated through diverse populations of spinal interneurons (Alstermark and Isa, 2012; Alstermark and Ogawa, 2004; Alstermark et al., 2004; Bernhard and Bohm, 1954; Fetz et al., 2002; Ueno et al., 2018; Yang and Lemon, 2003). Even in primates, only a minority of motor neurons receive direct cortical synaptic input, and this population is likely enriched in motor pools controlling fractionated finger movements, although there is evidence of its dispensability for skilled grasp (Alstermark and Isa, 2012; Alstermark et al., 2011; Porter and Lemon, 1993). Finally, our results show that corticospinal neurons extensively collateralize in many brain regions that in turn project densely to the spinal cord, and make notable contributions to patterning motor output. In light of this biological complexity, it is unsurprising that the motor cortical activity we observe is diverse and not exclusively linear in its relationship with motor output. Corticospinal neurons with activity abstractly related to muscle output may form collateral synapses in different brain regions or on different cell types than those with activity linearly related to muscle output. This may be further reflected in their spinal targets: those CSNs with activity more correlated with that of muscles might be more closely positioned to motor output, perhaps forming synapses on premotor interneurons like V2a subpopulations. Those with non-muscle-like activity may synapse on neurons implicated in gain control of proprioceptive sensory feedback or other modulatory functions. Future experiments using electrophysiological and transsynaptic mapping techniques – particularly those amenable to the spinal cord – will be invaluable in identifying these connectivity principles.

### Implications for the role of basal ganglia in sequencing behaviors

The basal ganglia play a critical role in the learning and performance of movement sequences (Agostino et al., 1992; Jin and Costa, 2015; Miyachi et al., 1997). For instance, perturbations of the dorsolateral striatum degrade the performance of learned lever press sequence behavior (Jin and Costa, 2010; Jin et al., 2014). Moreover, striatal SPNs encode features of movement sequences in their spiking activity, including a substantial population that encode lever press sequences as singular actions. While sensorimotor cortex is a major source of excitatory input to the striatum, the role motor cortex, and particularly CSNs, play in shaping striatal activity has been elusive. One reasonable possibility is that CSNs relay corollary discharge signals (i.e. efference copies of planned or ongoing movements) to the striatum, so that basal ganglia circuits have an accurate representation of actions (Alexander et al., 1986). Our results from experiments leveraging 2p imaging of CSNs combined with transsynaptic rabies tracing comport with this hypothesis, in that CSNs with identified striatal synapses display activity patterns indistinguishable from the broader CSN population. Interestingly, we found that equivalent information is encoded by CSNs_DLS_ synapsing on either the direct and indirect pathways of the basal ganglia, supporting the idea that CSNs act in a broadcasting capacity, leaving the translation of intent into action to downstream circuits in the basal ganglia and spinal cord (Arber and Costa, 2018). However, with any rabies-based tracing method, the absence of labeling does not indicate absence of connectivity, so our results likely undersample the abundance of CSNs with striatal synapses. In addition, rabies tracing methods do not reflect the strength of synaptic connectivity – a factor that likely determines the influence of CSN activity of distinct striatal circuits. To this end, our electrophysiological mapping experiments reveal a synaptic bias by which CSN activity could be overrepresented in the direct pathway of the basal ganglia through additional synapses on D1 SPNs. Moreover, dopaminergic feedback may enhance this dichotomy through opposing influences on D1 versus D2 SPN excitability (Albin et al., 1989; DeLong, 1990; Tritsch and Sabatini, 2012).

### Implications for the control of spinal motor output and sensory feedback

Rabies tracing experiments reveal that CSNs_DLS_ innervate broad regions of spinal cord, including both dorsal and ventral laminae of the spinal grey. Axon terminations are predictably the densest in cervical spinal cord, consistent with the role cervical motor pools play in controlling forelimb joints (Kandel et al., 2000). Yet, we observe substantial collateralization of CSNs that terminate in cervical levels, with axonal varicosities as far caudal as lumbar spinal cord (Leyton and Sherrington, 1917). Such an architecture may be useful in maintaining postural stability by increasing the excitability of motor pools innervating body wall muscles, many of which lie in thoracic spinal cord (Watson et al., 2009). Moreover, interneurons influence motor output through long range, intersegmental projections, raising the possibility that these thoracic and lumbar collaterals target interneurons that project back to cervical cord (Illert et al., 1977). Future experiments using electrophysiological circuit mapping and focal ablation methods will reveal the role these corollary synapses play in the coordination of intra-segmental spinal circuits.

We observed significant differences between the spatial distribution of CSN_D1_ and CSN_D2_ spinal projections, which possibly translates into differences in connectivity with subtypes of spinal interneurons. Indeed, recent studies have identified transcriptional profiles of functionally distinct spinal interneuron subgroups, many of which settle in restricted spatial compartments (Bikoff et al., 2016; Gabitto et al., 2016). One theory argues for the position of spinal neurons as a determinant of connectivity, begging the question of whether subgroups of CSNs form more connections with the subtypes of interneurons that settle in these compact regions (Balaskas et al., 2019; Surmeli et al., 2011). For instance, the density of synapses just dorsolateral to the central canal roughly overlaps with a large population of GAD2-expressing interneurons responsible for presynaptic inhibition of sensory afferents (Betley et al., 2009). Cortical projections to this population may be important for maintaining stable forelimb movements, given the outsized role GAD2-expressing interneurons play in preventing oscillatory limb movement by regulating proprioceptive feedback (Akay et al., 2014; Fink et al., 2014). This circuit is also well positioned to mediate a highly localized source of peripheral sensory filtration, where corollary discharge signals from motor cortex may engage inhibitory interneurons, which in turn could suppress the sensory consequences of movement (Crapse and Sommer, 2008). Future studies focused on the spinal consequences of CSN activity will be useful in exploring these possibilities.

We found that subgroups of CSNs that innervate one of multiple genetically-defined spinal interneurons also innervate DLS, furthering the possibility that synapse-specific subpopulations differentially influence striatal function. For instance, CSNs_Chx10_, CSNs_GAD2_, or CSNs_SST_ may preferentially synapse on either D1 or D2 SPNs, or on spatially-restricted clusters of SPNs. Because CSNs form more compact (Hooks et al., 2018) and less potent (Figures 2 and S4) terminations in striatum compared to IT corticostriatal neurons, a highly granular and specific coordination of spinal and striatal connectivity is feasible. However, substantially more research is needed to delineate the role of spinal interneuron subtypes in shaping motor output and sensory feedback. Extending this line of research will be important to understanding how corticospinal output shapes behavior through control of both spinal and striatal circuits.

Finally, we identified a major source of input to CSNs_DLS_ being thalamic nuclei that serve as output regions of the basal ganglia (Kuramoto et al., 2015). The notion that layer 5b neurons receive abundant thalamic input aligns with several reports that disrupt the orthodoxy of thalamic input being multiple synapses presynaptic to output of the canonical cortical column (Guo et al., 2018). Instead, our results indicate an anatomical basis by which the basal ganglia have unprecedented access to cortical control over spinal cord through synapses onto corticospinal neurons.

## Acknowledgements

We thank K. Fidelin and V. Athalye for feedback on this manuscript. We thank H. Rodrigues for designing and constructing behavioral equipment. We thank S. Brenner-Morton for custom antibodies, and S. Fageiry & K. Ritola for custom viral constructs. We are grateful for technical assistance from G. Martins, M. Correia, C. Warriner, A. Miri, and K. MacArthur. We thank I. Marcelo for time warping code. We thank T. Jessell for inspiring this research and for his invaluable critical feedback. R.M.C was funded by the National Institute of Health (5U19NS104649) and the Simons-Emory International Consortium on Motor Control. A.N. was a Howard Hughes Medical Institute Fellow of the Helen Hay Whitney Foundation and is currently supported by NIH Pathway to Independence Award 1K99NS118053-01.

## Author Contributions

A.N. and R.M.C designed experiments, interpreted data, and wrote this manuscript. A.N. performed experiments and analyzed data. B.A. assisted in collecting and analyzing anatomical tracing data.

## Methods

### Key Resource table information

| REAGENT or RESOURCE | SOURCE | IDENTIFIER |
| --- | --- | --- |
| **Antibodies** |  |  |
| Rabbit Anti-GFP | Columbia University | - |
| Living Colors® DsRed Polyclonal Antibody | Takara | 632496 |
| Guinea Pig Anti-vGlut1 | Columbia University | - |
| Rabbit Anti-Cre | MilliporeSigma | 69050-3 |
| Neurotrace 640/660 | ThermoFisher | N21483 |
| Alexa Fluor® 488 AffiniPure Alpaca Anti-Rabbit IgG (H+L) | Jackson ImmunoResearch | 611-545-215 |
| DyLight™ 405 AffiniPure Goat Anti-Guinea Pig IgG (H+L) | Jackson ImmunoResearch | 106-475-003 |
| Cy™3 AffiniPure Goat Anti-Rabbit IgG (H+L) | Jackson ImmunoResearch | 111-165-003 |
| Streptavidin, Alexa Fluor™ 405 conjugate | Thermo-Fisher | S32351 |
| **Virus Strains** |  |  |
| AAV1-CAG-FLEX-GFP | Penn Vector Core | CS0871 |
| AAV-retro-EF1α-Cre.mCherry | Addgene | 55632-AAVrg |
| AAV-retro-CAG-ChR2-tdTomato | Addgene | 28017-AAVrg |
| AAV-retro-CAG-GCaMP6f | Janelia Farm | - |
| AAV-2/1-Ef1α-FLEX-TVA.mCherry | UNC | AV5006B |
| AAV-2/1-CAG-FLEX-N2cG-mKate2.0 | Janelia Farm | - |
| AAV-retro-Ef1α-fDIO-Cre | Addgene | 121675-AAVrg |
| AAV-retro-Ef1α-FlpO | Addgene | 55637-AAVrg |
| AAV-DJ-CAG-FRT-synaptophysinGFP | Janelia Farm | - |
| AAV-retro-CAG-FLEX-GFP | Addgene | 51502-AAVrg |
| EnVA-N2cΔG-tdTomato | Janelia Farm | - |
| AAV-DJ-fDIO-eYFP | UNC | AV6220C |
| EnVA-N2cΔG-FlpO.mCherry | Janelia Farm | - |
| **Chemicals, Peptides, and Recombinant Proteins** |  |  |
| Strontium Chloride Hexahydrate | Sigma-Aldrich | 255521 |
| DAPI | Thermo-Fisher | D1306 |
| TrueBlack | Biotium | 23007 |
| **Experimental Models: Organisms/Strains** |  |  |
| Mouse: C57BL/6J | The Jackson Laboratory | 000664 |
| D1-tdTomato | The Jackson Laboratory | 016204 |
| D2-GFP | MMRRC | 000230 |
| D1-Cre (EY217) | MMRRC | 030778 |
| A2a-Cre | MMRRC | 036158 |
| Chx10-Cre | Custom | - |
| GAD2-Cre | The Jackson Laboratory | 010802 |
| SST-Cre | The Jackson Laboratory | 028864 |
| **Software and Algorithms** |  |  |
| MATLAB 2018a | Mathworks |  |
| Imaris | Bitplane |  |
| ImageJ | NIH |  |
| **Other** |  |  |
| Two-photon microscope and data acquisition system | Bruker | - |
| 4-channel EMG amplifier | University of Cologne<br>Electronics Lab | MA 102S |
| pyControl | https://pycontrol.readthedocs.io/en/latest/ |  |
| Stainless steel wire for EMG electrodes | A-M Systems | 793200 |

### EXPERIMENTAL MODEL AND SUBJECT DETAILS

All experiments and procedures were performed according to NIH guidelines and approved by the Institutional Animal Care and Use Committee of Columbia University.

#### Experimental Animals

Adult mice of both sexes, aged between 2-6 months were used for all experiments, including slice electrophysiology. The strains used were: C57BL6/J, Jackson Laboratories #000664, B6.Cg-Tg(Drd1a-tdTomato)6Calak/J, Jackson Laboratories #016204, Tg(Drd2-EGFP)S118Gsat/Mmnc, MMRRC #000230, Tg(Drd1a-cre)EY217Gsat/Mmucd, Jackson Laboratories #030778, B6.FVB(Cg)-Tg(Adora2a-cre)KG139Gsat/Mmucd, MMRRC #036158, Chx10-Cre, Custom Jessell Laboratory, B6J.Cg-Ssttm2.1(cre)Zjh/MwarJ, Jackson Laboratories #028864, Gad2tm2(cre)Zjh/J, Jackson Laboratories #010802. Mice used for behavioral experiments were individually housed, and all mice were kept under a 12 hour light/dark cycle.

## Methods Detail

### Stereotaxic Viral Injections

Analgesia in the form of subcutaneous injection of carprofen (5 mg/kg) or buprenorphine SR (0.5-1mg/kg) was administered the day of the surgery, along with bupivacaine (2mg/kg). Mice were anesthetized with isoflurane and placed in a stereotaxic holder (Leica). A midline incision was made to expose the skull, and a craniotomy was made over the injection site. To label CSNs with GFP, AAV-FLEX-GFP, 100nL of virus was injected into each of two sites of motor cortex, 1.5mm lateral to the midline and 0.5 and 1.0mm rostral to bregma, approximately 700μm below the pial surface. Care was made to ensure there was no efflux of virus by stabilizing the skull and waiting 10 minutes after penetration before injecting. AAV-retro-Cre.mCherry was then injected into spinal cord (see below). For rabies-based transsynaptic tracing from striatal SPNs, AAV-FLEX-N2cG and AAV-FLEX-TVA.mCherry, 40nL of a 1:1 mixture was injected into DLS at 0.5mm rostral, 2.65mm lateral, and 3.5mm ventral to bregma. To label CSN axons in spinal cord, a vertical approach was taken to target DLS, and 300nL of EnVA-N2cΔG-tdTomato was injected. For transsynaptic tracing following 2p imaging, a craniotomy was made just caudal to the cranial window. The injection pipette was angled along the rostrocaudal axis, and the same region of DLS targeted for injections of rabies helper viruses was injected with 300nL of pseudotyped deficient rabies virus. To express ChR2 in intratelencephalic neurons, 100nL of AAV-retro-ChR2.tdTomato was injected into either motor cortex or DLS contralateral to the hemisphere targeted for whole cell recording. For transsynaptic rabies tracing experiments to label inputs to CSNs_DLS_, 100nL of AAV-retro-FRT-Cre was injected into DLS, AAV-retro-FlpO was injected into spinal cord (see below), and 100nL of a 1:1 mixture of AAV-FLEX-N2cG and AAV-FLEX-TVA.mCherry was injected into forelimb motor cortex. Two weeks later, 300nL of EnVA-N2cΔG-tdTomato was injected into motor cortex. To tag CSNs_DLS_ with GFP, AAV-retro-FLEX-GFP was injected into DLS, and AAV-retro-Cre.mCherry was injected into spinal cord (see below). To tag subpopulations of CSNs, 250nL of AAV-FRT-EYFP was injected into forelimb motor cortex at each of two sites. AAV-FLEX-N2cG and AAV-FLEX-TVA.mCherry was then injected into spinal cord, followed two weeks later by injections of EnVA-N2cΔG-FlpO.mCherry into spinal cord (see below).

### Spinal Cord Viral Injections

Analgesia in the form of subcutaneous injection of carprofen (5 mg/kg) or buprenorphine SR (0.5-1mg/kg) was administered the day of the surgery, along with bupivacaine (2mg/kg). Mice were anesthetized with isoflurane and placed in a stereotaxic holder (Leica). A midline incision was made to expose the spinal column. The muscular overlying the column was resected, and a metal clip attached to a spinal clamp was used to secure the T2 process and minimize spinal cord movement. The tail was gently stretched with another spinal clamp and separate the vertebrae. A surgical microknife and fine forceps were used to sever the meninges, exposing the spinal cord. A pulled glass pipette was filled with virus, and a Nanoject III was used to make multiple small volume injections across into the spinal cord, with parameters that depended on the experiment and reagents used. For injections of AAV-retro-GCaMP6f, AAV-retro-Cre.mCherry, AAV-retro-ChR2.tdTomato, or AAV-retro-FlpO, one penetration was made into each segment of the spinal cord between C3 and C8. Twenty injections of 10nL each were made into the center of the spinal grey, for a total volume of 200nL per spinal segment. For injections of AAV-FLEX-N2cG or AAV-FLEX-TVA.mCherry, two penetrations were made into each segment of the spinal cord between C3 and C8. 10nL of virus was injected along the dorsoventral axis every 50μm between 1.2mm and 0.1mm below the surface of the cord, totaling 480nL per segment. For injections of EnVA-N2cΔG-FlpO.mCherry, three penetrations were made into each segment of the spinal cord between C3 and C8. 15nL of virus was injected along the dorsoventral axis every 50μm between 1.2mm and 0.1mm below the surface of the cord, totaling 1080nL per segment. Following all injections, the skin was sutured closed and animals were closely monitored during recovery.

### Slice Electrophysiology and Optogenetic Photostimulation

Mice were deeply anesthetized with isoflurane and transcardially perfused with an ice-cold carbogenated high magnesium (10mM) ACSF. The brain was removed from the skull, and glued to the stage of a vibrating microtome (Leica). 300μm coronal brain slices were cut in a bath of ice-cold, slushy, carbogenated low calcium ACSF. Slices were incubated for 15-30 minutes in a 37°C bath of normal ACSF containing (in mM): 124 NaCl, 2.7 KCl, 2 CaCl2, 1.3 MgSO4, 26 NaHCO3, 1.25 NaH2PO4, 18 glucose, 0.79 sodium ascorbate. Slices were then transitioned to room temperature, where they remained for the duration of the experiment. Patch electrodes (3-6MΩ) were filled with either a potassium gluconate based internal solution (135 mM K-gluconate, 2 mM MgCl2, 0.5 mM EGTA, 2 mM MgATP, 0.5 mM NaGTP, 10 mM HEPES, 10 mM phosphocreatine, 0.15% Neurobiotin) or a cesium/QX-314 based internal solution (5 mM QX-314, 2 mM ATP Mg salt, 0.3 mM GTP Na salt, 10 mM phosphocreatine, 0.2 mM EGTA, 2 mM MgCl2, 5 mM NaCl, 10 mM HEPES, 120 mM cesium methanesulfonate, and 0.15% Neurobiotin). All recordings were made using a Multiclamp 700B amplifier, the output of which was digitized at 10 kHz (Digidata 1440A). Series resistance was always <35 MΩ and was compensated up to 90%. Neurons were targeted with differential interference contrast (DIC) and epifluorescence when appropriate. For simultaneous recordings, pairs of neighboring SPNs (within 50 μm of each other) were identified first by morphology using DIC imaging. The cellular identity of targeted neurons was confirmed through expression or lack of expression of transgenically-targeted fluorescent reporters. For experiments exploiting potassium gluconate based internal solutions, neurons were further identified through intrinsic electrophysiological properties, including excitability and current/voltage transformation. In a subset of experiments, cell morphology was visualized through internal dialysis of 0.1 mM Alexa Fluor 594 cadaverine or 0.1 mM Alexa Fluor 488 Na salt. ChR2-expressing axons were photostimulated using 10ms pulses of 473nm LED light (CoolLED) delivered through a 10x objective centered over the recording site. Brain slices were histologically processed to visualize Neurobiotin-filled cells through streptavidin-Alexa Fluor processing.

### Histology and Confocal Imaging

Mice were deeply anesthetized with isoflurane and transcardially perfused with phosphate buffered saline (PBS) followed by ice cold 4% paraformaldehyde. Brains and spinal cords were post-fixed overnight in 4% paraformaldehyde, and then cryopreserved in a 30% sucrose solution for 3 days at 4°C. Brains and spinal cords were embedded in Optimum Cutting Temperature Compound (Tissue-Tek), and 70μm coronal sections were cut on a cryostat. Tissue was rinsed several times in PBS, then permeabilized in PBS containing 0.2% Triton X-100 (PBST). For imaging synapses in spinal cord, tissue sections were first permeabilized in 1% PBST to aid in antibody penetration. Immunostaining was performed with primary antibodies diluted at 1:1000 for 3 days at 4°C, and with secondary antibodies at 1:000 overnight at 4°C. Counterstains of DAPI or Neurotrace were included in the secondary antibody incubation at 1:1000. For imaging synapses, brain slices mounted to slides were briefly incubated with TrueBlack diluted in 70% ethanol to quench lipofuscin and background autofluorescence. Confocal imaging was performed on a Zeiss 710 or Zeiss 880 using 10x, 20x, 40x, 63x, or 100x objectives. For mapping the distribution of spinal synapses arising from CSNs_DLS_, high XYZ resolution stitched images were acquired overnight using a 40x water immersion high NA objective. Imaris was used to identify tdTomato^+^ axons that colocalize with vGlut1 expression. Synaptic boutons were then marked with spots, and the coordinates of these spots were measured relative to the center of the central canal.

### Slide Scanning and Anatomical Reconstructions

70μm coronal sections were serially mounted on slides, and were treated with TrueBlack diluted in 70% ethanol to quench lipofuscin and background autofluorescence. Sections were imaged using an AZ100 automated slide scanning microscope equipped with a 4X 0.4NA objective. (Nikon). Image processing and analysis using BrainJ proceeded as previously described (Botta et al., 2019). Briefly, brain sections were aligned and registered using 2D rigid body registration. A machine learning pixel classification approach using Ilastik was employed to identify cell bodies and neuronal processes. To map the location of these structures to an annotated brain atlas, 3D image registration was performed using Elastix relative to a reference brain. The coordinates of detected cells and processes were then projected into the Allen Brain Atlas Common Coordinate Framework. Visualizations of the data were performed in ImageJ and Imaris, and subsequent analyses were performed in MATLAB using custom software.

### Electromyographic Electrode and Headpost Implantation

Electromyographic electrodes were fabricated as previously described (Akay et al., 2006). Two pieces of insulated braided stainless-steel wire were knotted, and half-millimeter portions of insulation were stripped from each wire just below the knot, so that exposed contact sites were separated by 0.5 millimeter. The portions of wire with contact sites were twisted, and the ends secured in a crimped hypodermic needle to permit easy insertion into targeted muscle groups. The opposing strands were soldered to a miniature connector. This process was repeated three times to produce a total of four differential recording electrodes that could be implanted into four muscles.

Analgesia in the form of subcutaneous injection of carprofen (5 mg/kg) or buprenorphine SR (0.5-1mg/kg) was administered the day of the surgery, along with bupivacaine (2mg/kg). Mice were anesthetized with isoflurane and placed in a stereotaxic holder (Leica). Hair was carefully shaved from the right forelimb, neck, and head, and the skin was thoroughly cleaned. Incisions were made over the neck and forelimb, and the electrode assemblage was snaked through these sites so that the miniature connector was positioned near the head and the individual recording electrodes positioned near biceps, triceps, extensor digitorum communis, and palmaris longus. Electrodes were implanted in each muscle by passing each needle and wire through targeted muscle groups until the knot was abutted to the muscle entry point. The tag ends of wire were then knotted by the exit point, thus securing the contact sites within the muscle. Forelimb incisions were closed with sutures, and the headpost implantation proceeded. The scalp was removed to expose the cranium, and facia was cleared using a scalpel and saline irrigation. A custom, 3D printed plastic headpost was affixed to the cranium using Metabond dental cement (Parkell), and reference points were marked to facilitate the implantation of a cranial window. Finally, the miniature connecter for the EMG electrode assemblage was cemented to the caudal edge of the headpost, and the skin overlying the neck was closed with sutures.

### Cranial Window Implantation

Analgesia in the form of subcutaneous injection of buprenorphine SR (0.5-1mg/kg) and was administered the day of the surgery, along with bupivacaine (2mg/kg) and the anti-inflammatory dexamethasone (2mg/kg). Mice were secured in a stereotaxic frame (Leica) and the head was secured using 3D printed forks designed to clamp the custom headpost. The custom cranial window was composed of two semicircular pieces of glass coverslip (200 μm thick, Tower Optical Corp.), fused together and then to a 4mm round #1 coverslip (Warner Instruments) with optical cement (Norland Optical Adhesive 61). A craniotomy the shape of the insertable coverslips was made over forelimb motor cortex, and the window was implanted so that the semicircular plug was gently pressing on the brain. The entire assemblage was secured using Metabond.

### Behavior

Behavioral training occurred in parallel using behavioral chambers equipped with custom-made and assembled components. Mice were head-fixed using 3D printed hard plastic forks that clamped around a custom plastic headpost cemented to the cranium. The body rested in an opaque plastic tube, and the left forelimb was allowed to rest on a moveable perch. The right forelimb was positioned over a milled plastic lever that had a small counterweight. Lever presses were reported as the counterweighted arm passed through an infrared beam. Water rewards were dispensed through a blunt needle positioned ~3mm from the mouth so that beads of water reward were reachable by licking. Water reward was calibrated regularly by adjusting the length of the TTL pulse sent to a solenoid valve. Behavioral assays were controlled using software written for and deployed with pyControl (https://pycontrol.readthedocs.io/en/latest/). Performance was continuously monitored and recorded with webcams.

Mice were accustomed to handling for several days, and then placed on a water restriction schedule using established guidelines (Guo et al., 2014). Weight, appearance, and general health was monitored daily, and supplemental water was administered when necessary. Water-restricted mice were acclimated to the custom-made behavioral apparatus for two days, where they received water reward (5μL) at sporadic intervals for 15 minutes (day 1) or 30 minutes (day 2). For the first phase of training (~10 days), mice were required only to press the lever once to receive reward. A timeout period of 3 seconds following reward was imposed to discourage continuous pressing, and the session ended when reward volume totaled 1000μL or one hour passed. Supplemental water was given to ensure an adequate daily volume. For the second phase of training (~7 days), reward was delivered after every forth lever press, regardless of the inter-press interval duration. For the final phase of training (~14 days), a countdown was imposed requiring four lever presses to occur within 2 seconds in order to receive reward. The countdown was reset after reward delivery, and a 3 second timeout was imposed.

### Two-photon Imaging

Calcium imaging experiments were performed using a modified two-photon microscope (Bruker) outfitted with a 25x 1.0NA water immersion objective (Olympus) and a mode locked Ti:sapphire laser (Verdi 18W, Coherent) at 940nm. A custom-made computerized, motorized goniometer was used to subtly and reproducibly angle the head so that the cranial window was orthogonal to the beam path. Images were acquired using Prairie View software (Bruker) at 64Hz, and every 4 images were averaged, yielding an effective sampling rate of 16Hz. Data was acquired from an area approximately 430μm x 430μm with 256 x 256 pixels. Multiple non-overlapping field of view were imaged from each mouse over ~7 days. Following injections of EnVA-N2cΔG-tdTomato, fields of view from functional imaging sessions were identified by first aligning surface vasculature, then carefully aligning basal GCaMP fluorescence signals to reference images taken during functional imaging. Z stacks and 2D images of tdTomato fluorescence were acquired at a wavelength of 1040nm.

### Electromyographic Recordings

EMG signals were amplified and filtered (250-20,000 Hz) with a differential amplifier (MA102 with MA103S preamplifiers, University of Cologne electronics lab). These signals were acquired at 10kHz alongside two-photon imaging data using Prairie View. EMG signals were down-sampled to 1kHz, high-pass filtered at 40Hz, rectified, and convolved with a Gaussian that had 10 ms standard deviation.

### Quantification and Statistical Analyses

#### Automated Anatomical Reconstruction

Analysis of slide scanning data was performed using MATLAB. Data was output from the BrainJ pipeline in the form of CSV files containing measurements of neurite labeling and cell body count from each region in the Allen Brain Atlas Common Coordinate Framework. These measurements were hierarchically organized so that analyses from sub-regions (i.e. layers of primary motor cortex) could be performed alongside more general annotations (i.e. primary motor cortex). For measurements from high order ancestor regions (i.e. isocortex), measurements from descendent regions identified by Allen Brain Atlas application programming interface were grouped.

#### Slice Electrophysiology

Analysis of slice electrophysiology data was performed in MATLAB and in Clampfit (Molecular Devices). Tests of significance were performed using paired t-tests with an alpha of 0.05. Amplitude and charge were measured from a 200ms window following stimulus onset relative to a baseline period 250ms before the onset of stimulus. To measure the amplitude of miniature EPSCs evoked through optogenetic stimulation of CSNs using strontium-containing ACSF, a mEPSC template was created in Clampfit. That template was used to search for mEPSCs in the tail response following the early synchronous release of neurotransmitter. Each mEPSC was manually reviewed, misidentified events were excluded from analysis, and the resulting mEPSCs were averaged for each cell.

#### Calcium Imaging

Calcium imaging analysis was performed using constrained non-negative matrix factorization (CNMF). First, raw imaging datasets (~10 minutes each) were motion corrected using rigid, then non-rigid registration. Registered datasets were then processed in CNMF using an autoregressive process *p* of 2. Analysis was also performed using a p of 0 to replicate results, although this data is not included in this study. Output of the CNMF was in the form of ΔF/F and inferred spike rate. Signals were up-sampled to match the sampling rate of EMG data, and Z-scored for further analysis. Lever press trials were time warped by expanding or contracting inter-press intervals using linear resampling to match a template with fixed intervals of 200ms. Neurons were classified by their response properties as follows. For each neuron, trials of 4 lever press sequences were identified and time warped. A baseline period was defined as the first 250ms of each trial (beginning 1.5sec before the first lever press). Each trial (excluding the baseline period) was segmented into bins 10 samples in length, and the bins with mean activity significantly different than baseline (measured using within-trial paired t-test) were marked. Within this group, bins with mean activity greater than 2.5 standard deviations of baseline were then identified as positively modulated, and bins with mean activity less than that of baseline were identified as negatively modulated. We then identified significantly modulated bins in each of 9 time periods that spanned the trial (excluding the baseline period). The rationale for analyzing short bins was that brief deviations in activity could be overlooked or diluted if averaging across longer time windows. Neurons with significant and positively modulated activity in one or more of periods 1-4, and zero in periods 6-9 were classified as ON. Neurons with significant and positively modulated activity in one or more of periods of 6-9, and zero in periods 1:4 were classified as OFF. Neurons with significant and positively modulated activity in two or more of periods 3-7 were marked as SUS. Neurons with significant and negatively modulated activity in two or more of periods 3:7 were classified as SUPR. Neurons that met none of these criteria were marked as UN.

To mark CSNs co-labeled with tdTomato through rabies infection, we used a 3D reconstruction approach to improve identification of red fluorescent neurites. Around one week to ten days after rabies injection, Z stacks of tdTomato fluorescence were acquired at 1040nm. These tdTomato Z stacks were imported into Imaris, and binary masks were generated using the surfaces function. The binary stack was then resliced to generate one binary mask at the same Z plane used for functional imaging of GCaMP. This mask was registered to the functional data set using shift parameters derived from registration of reference GCaMP fluorescence images. We then identified tdTomato pixels that fell within the spatial boundaries of GCaMP ROIs, and summed these pixels, which were weighted by how close they were to the center of the ROI. This number was divided by the total tdTomato pixels within that structure, yielding a value that reflected 1) the proximity of the tdTomato process to the center of the GCaMP ROI, and 2) the degree to which the tdTomato structure was overlapping with the GCaMP ROI. If this value was greater than 60% of the sum of weighted GCaMP ROI pixels divided by the total number of those pixels, that ROI was marked as tdTomato^+^.

## Supplementary Information

**Figure 1S. Mapping brainwide inputs to the spinal cord**

(A) Illustration of experimental approach to visualize cellular inputs to cervical spinal cord. (B) 3D reconstruction of brainwide inputs to spinal cord. Colors correspond to major brain divisions in which they reside. (C) The top brain regions that provide input to spinal cord, determined by the relative fraction of total identified somata. Notable brain regions are indicated by colored bars. Photomicrograph insets illustrate exemplar brain regions with substantial labeling. Dashed boxes are colored to correspond to notable brain regions from the bar graph. The inset pie chart shows the major ancestor brain structures projecting to cervical spinal cord. (D) Quantification of cortical inputs to spinal cord, divided by cortical region and laminae. The inset photomicrograph illustrates the L5b positioning of corticospinal neurons. Note that the Allen Brain Atlas classification did not subdivide L5 into L5a and L5b, and the position of corticospinal somata fell around the boundary between L5 and L6a. (E) Illustration of experimental strategy, same as Figure 1A. (F) Major ancestor brain regions containing GFP^+^ neurite. Note that this includes dendritic processes in sensorimotor cortex. Grouping the many brain regions comprising these ancestor structures reveals the intense innervation of several subcortical structures. (G) Experimental strategy to label synapses arising from CSNs. (H) Synaptophysin GFP (green) labeling in the brain. (I) Synaptophysin GFP (green) and FlpO (red) labeling in motor cortex. (J) Synaptophysin GFP (green) labeling in DLS. (K) Top brain regions to which CSNs project, measured as what fraction of all synapses are found within those brain structures, excluding sensorimotor cortex and fiber tracts.

**Figure 2S. Mapping the brainwide targets of CSNs_DLS_**

(A) Experimental strategy to label corticospinal neurons that project to striatum (CSNs_DLS_). (B) Photomicrograph of CSNs_DLS_ and their projections to DLS. (C) 3D reconstruction of CSNs_DLS_ projections throughout the brain, colored by targeted brain region. (D) Quantification of cortical regions contributing to the total population of CSNs_DLS_, compared to experiments from Figure 1 targeting the motor cortical population of CSNs. (E) Quantification of brain regions targeted by CSNs_DLS_, compared to data from Figure 1. Note that – despite the differences in experimental strategy – DLS is a primary target of CSNs_DLS_.

**Figure 3S. Identifying neurons presynaptic to CSNs_DLS_**

(A) Strategy to use intersectional transsynaptic tracing to label inputs to CSNs_DLS_. (B-D) Identification of starter cells (arrowheads) through coexpression of tdTomato (B) and Cre (C). Overlay in (D). (E) 3D distribution of tdTomato^+^ neurons, color coded by brain group. (F-O) Example confocal micrographs of tdTomato labeling throughout the brain. DAPI is blue. (P) Quantification of the top brain regions giving rise to neurons that form synapses on CSNs_DLS_. The pie chart indicates the major brain groups providing input to CSNs_DLS_. The inset image displays neuronal labeling in subdivisions of the thalamus.

**Figure 4S. Synaptic organization of intratelencephalic corticostriatal projections**

(A) Schematic illustrating the experimental strategy. Retrogradely-transported and expressed AAV encoding ChR2.tdTomato was injected into contralateral DLS or M1. D1 and D2 SPNs were targeted for simultaneous recording. (B) Photomicrograph of ChR2.tdTomato (red) and D2-GFP (green) labeling in a brain slice. (C) High magnification image of the boxed region from (B). Note the expansive axonal plexus. (D) DIC image of a D1^+^ (magenta) and D1^−^ SPN targeted for simultaneous whole cell recording. The dashed lines indicate the location of recording electrodes. (E) Superimposed current-clamp voltage recordings from an SPN following optogenetic stimulation of IT corticostriatal axons, highlighting the potency of this projection. (F) Grand average response of all D1 (blue) and D2 (orange) SPNs to optogenetic stimulation of IT corticostriatal neurons. (G-H) Pairwise comparison of ChR2-evoked amplitude (G) and charge (H) in D1 versus D2 SPNs. (I-J) Trial average of mEPSC evoked from an example D1 (I) and D2 (J) SPN. Individual trials are in grey. (K) Average mESPC amplitudes in D1 versus D2 SPNs. (L) Distribution of all mEPSCs ordered by mEPSC peak current, recorded in D1 (blue) or D2 (orange) SPNs. The inset is an overlay of the average mEPSC from D1 and D2 SPNs.

**Figure 5S. Method to analyze the distribution of spinal synapses arising from CSNs_DLS_**

(A) Raw CSN spinal synapse data from three example mice. The position of each dot corresponds to a vGlut1^+^ axonal varicosity. (B) Raw data is spatially binned for each mouse. The A sliding window is used to group local bins, and the density of labeling within these groups is compared across genotypes of mice.

**Figure 6S. Mapping brainwide targets of CSNs defined by their spinal cellular targets.**

(A) Experimental strategy to drive expression of GFP in corticospinal neurons that form synapses on identified spinal cell types. (B) Cortical regions that contain CSNs that project to different spinal cell types. (C) Reliability of cell body labeling within and across genotypes. (D) Quantification of brain structures that receive substantial input from CSN subtypes. The inset is a 3D reconstruction of axons from CSNs_Chx10_, color coded by brain region.

**Figure 7S. Temporal heterogeneity of CSN activity around lever press**

(A) Z scored activity of corticospinal neurons aligned to single lever press events. (B) Histogram of the times of peak activity relative to lever press, for all neurons.

**Figure 8S. Analysis of CSN activity during lever press sequences**

(A-B) Illustration of time warping procedure for four press sequences. Dots indicate lever press times, as well as timepoint used for pre- and post-trial alignment (six time points per trial). (C-D) Z scored calcium activity before (C) and after (D) time warping. Note the emergence in (D) of peaks in activity corresponding to individual lever press events. (E) Plot of the top three principle components of normalized CSN activity.

**Figure 9S. Classification of CSN activity profiles**

(A) Histogram of the times of peak activity for CSNs with categorized activity profiles, aligned to lever press sequence onset.

**Figure 10S. Analysis of EMG during behavior and CSN activity correlations to EMG**

(A) Average EMG activity for four forelimb muscles aligned to single lever presses. (B) Average EMG activity for four forelimb muscles aligned to lever press sequences. (C) Correlation of CSN activity to biceps versus triceps EMG during concatenated random segments of behavior and rest (session, grey) or concatenated lever press sequences (purple), matched in duration. (D) Correlation of trial-averaged CSN activity with biceps or triceps EMG. Neurons with correlations biased to triceps or biceps are colorized in red or green, respectively. (E) Average lever press sequence-related activity of CSNs highly correlated to triceps (red) or biceps (green) EMG. Activity from neurons with similar correlation coefficients is in grey.

**Figure 11S. Method to identify CSNs with identified striatal synapses**

(A) Exemplar photomicrograph of CSNs expressing GCaMP (green), and corticostriatal neurons marked with tdTomato (red). (B) Cartoon depiction of fluorescent expression possibilities, viewed from an X-Z perspective. (C) Cartoon depiction of fluorescent expression possibilities, viewed from an X-Y perspective. (D) Two example possibilities for overlapping green and red fluorescence, one constituting a double-positive (top) and one rejected from being a double-positive (bottom).

