## Supplemental Figures for "Corticospinal neurons encode complex motor signals that are broadcast to dichotomous striatal circuits"

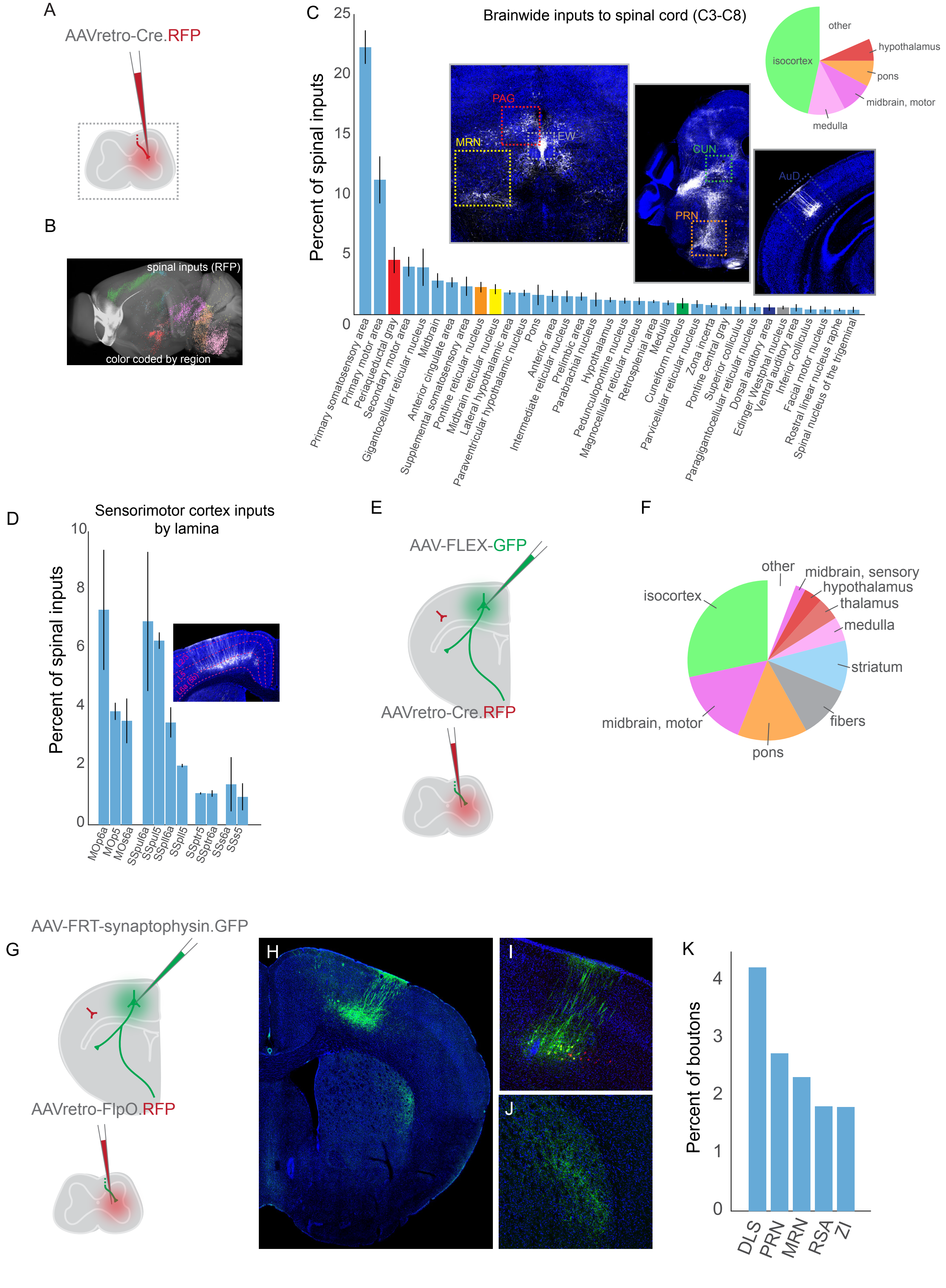
A
AAVretro-Cre.RFP
B
spinal inputs (RFP)
color coded by region
C
Brainwide inputs to spinal cord (C3-C8)
Percent of spinal inputs
PAG
EW
MRN
CUN
PRN
AuD
other
hypothalamus
isocortex
pons
midbrain, motor
medulla
Primary somatosensory area
Primary motor area
Periaqueductal gray
Secondary motor area
Gigantocellular reticular nucleus
Midbrain
Anterior cingulate area
Supplemental somatosensory area
Pontine reticular nucleus
Midbrain reticular nucleus
Lateral hypothalamic area
Paraventricular hypothalamic nucleus
Pons
Anterior area
Intermediate reticular nucleus
Prelimbic area
Parabrachial nucleus
Hypothalamus
Pedunculopontine nucleus
Magnocellular reticular nucleus
Retrosplenial area
Medulla
Cuneiform nucleus
Parvicellular reticular nucleus
Zona incerta
Pontine central gray
Superior colliculus
Paragigantocellular reticular nucleus
Dorsal auditory area
Edinger Westphal nucleus
Ventral auditory area
Inferior colliculus
Facial motor nucleus
Rostral linear nucleus raphe
Spinal nucleus of the trigeminal
D
Sensorimotor cortex inputs by lamina
Percent of spinal inputs
MOp6a
MOp5
MOs6a
SSpul6a
SSpul5
SSpll6a
SSpll5
SSptr5
SSptr6a
SSs6a
SSs5
E
AAV-FLEX-GFP
AAVretro-Cre.RFP
F
other
midbrain, sensory
hypothalamus
thalamus
medulla
striatum
fibers
pons
midbrain, motor
isocortex
G
AAV-FRT-synaptophysin.GFP
AAVretro-FlpO.RFP
H
I
J
K
Percent of boutons
DLS
PRN
MRN
RSA
ZI

Figure S1

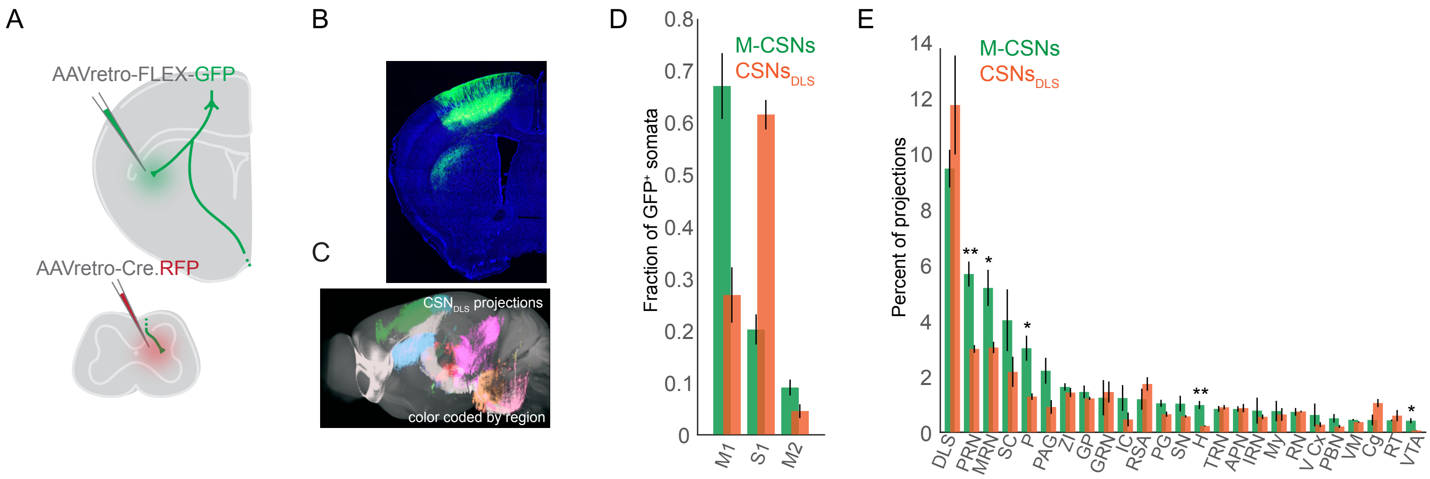

A
AAVretro-FLEX-GFP
AAVretro-Cre.RFP
B
C
CSNDLS projections
color coded by region
D
M-CSNs
CSNsDLS
Fraction of GFP+ somata
0
0.1
0.2
0.3
0.4
0.5
0.6
0.7
0.8
M1
S1
M2
E
M-CSNs
CSNsDLS
Percent of projections
0
2
4
6
8
10
12
14
**
*
*
**
*
DLS PRN MRN SC P PAG ZI GP GRN IC RSA PG SN H TRN APN IRN My RN V Cx PBN VM Cg RT VTA


### Figure S2

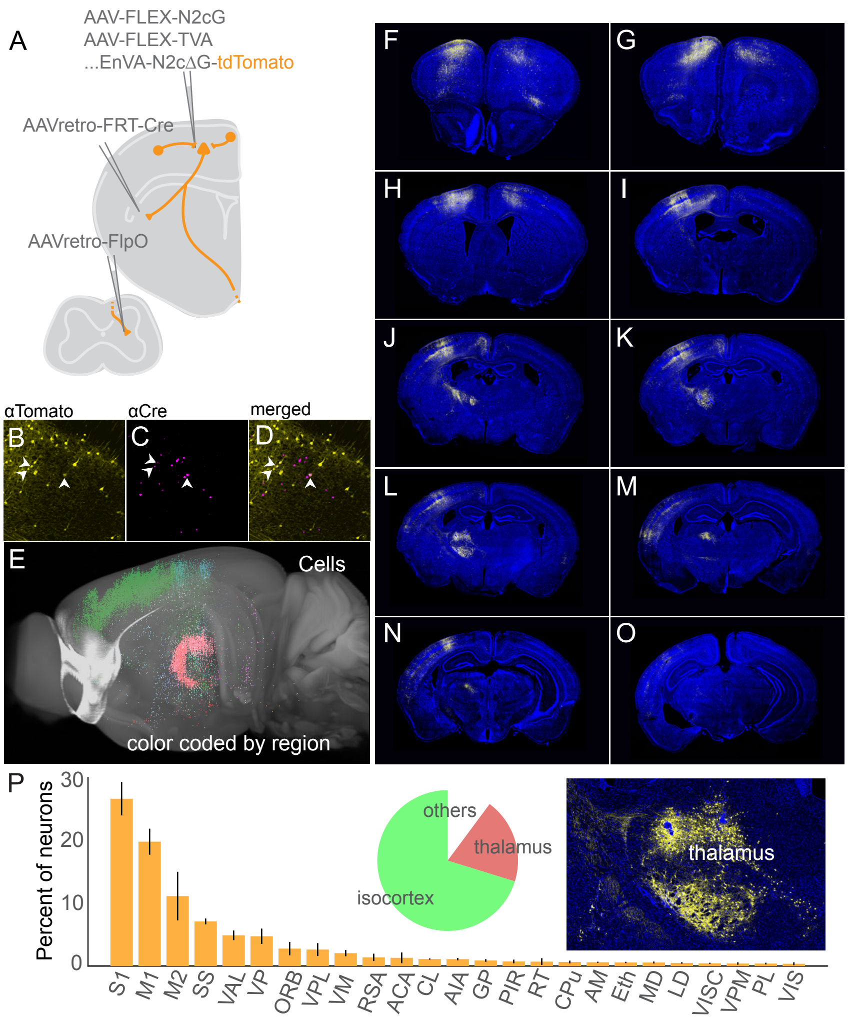
A
AAV-FLEX-N2cG
AAV-FLEX-TVA
...EnVA-N2cΔG-tdTomato
AAVretro-FRT-Cre
AAVretro-FlpO
αTomato
αCre
merged
B
C
D
E
Cells
color coded by region
F
G
H
I
J
K
L
M
N
O
P
Percent of neurons
30
20
10
0
S1
M1
M2
SS
VAL
VP
ORB
VPL
VM
RSA
ACA
CL
AIA
GP
PIR
RT
CPu
AM
Eth
MD
LD
VISC
VPM
PL
VIS
others
thalamus
isocortex
thalamus

Figure S3

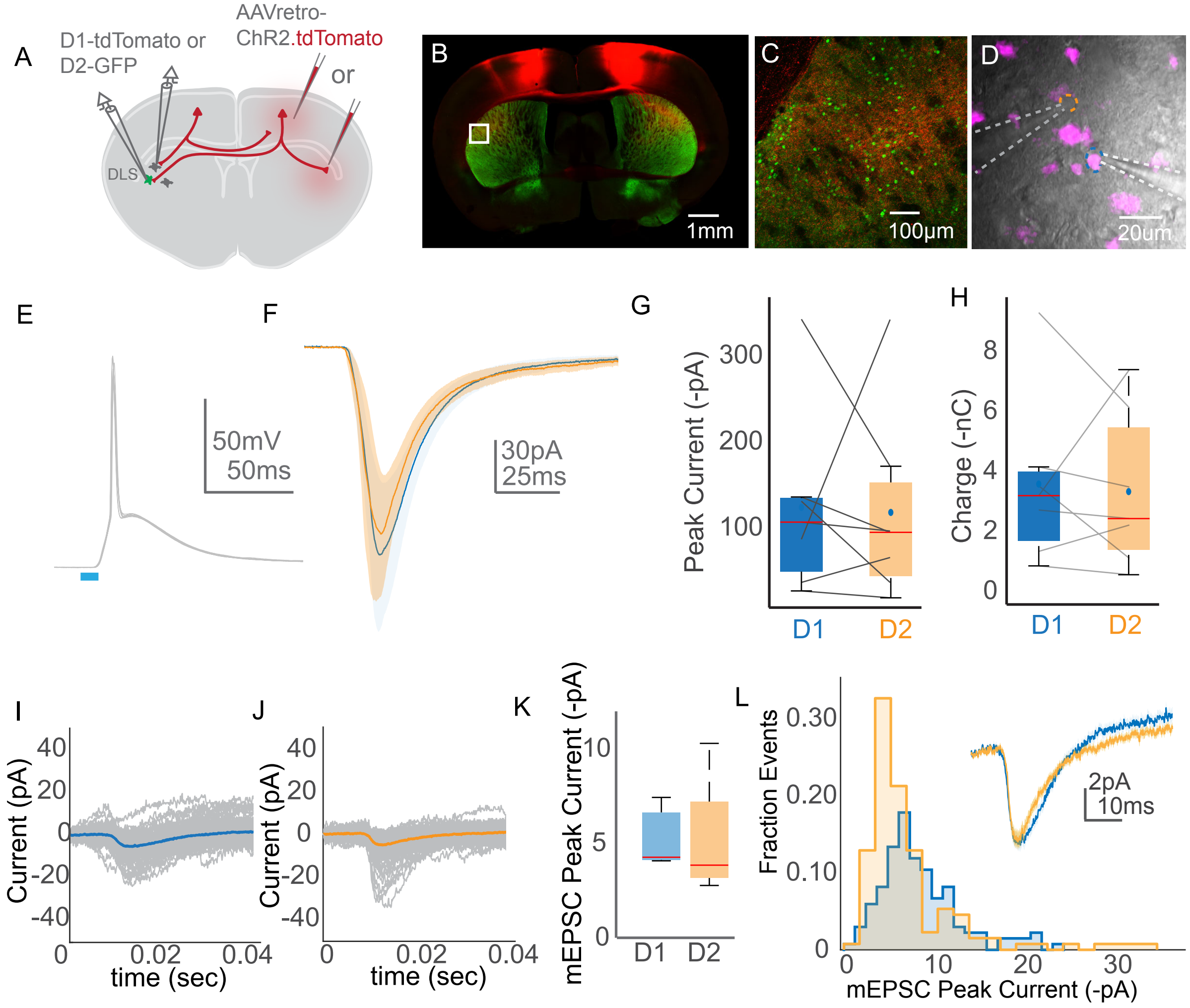
A
D1-tdTomato or
D2-GFP
AAVretro-
ChR2.tdTomato
or
DLS
B
1mm
C
100µm
D
20µm
E
50mV
50ms
F
30pA
25ms
G
Peak Current (-pA)
300
200
100
D1
D2
H
Charge (-nC)
8
6
4
2
0
D1
D2
I
Current (pA)
40
20
0
-20
-40
0
0.02
0.04
time (sec)
J
Current (pA)
40
20
0
-20
-40
0
0.02
0.04
time (sec)
K
mEPSC Peak Current (-pA)
10
5
0
D1
D2
L
Fraction Events
0.30
0.20
0.10
0
0
10
20
30
mEPSC Peak Current (-pA)
2pA
10ms

Figure S4

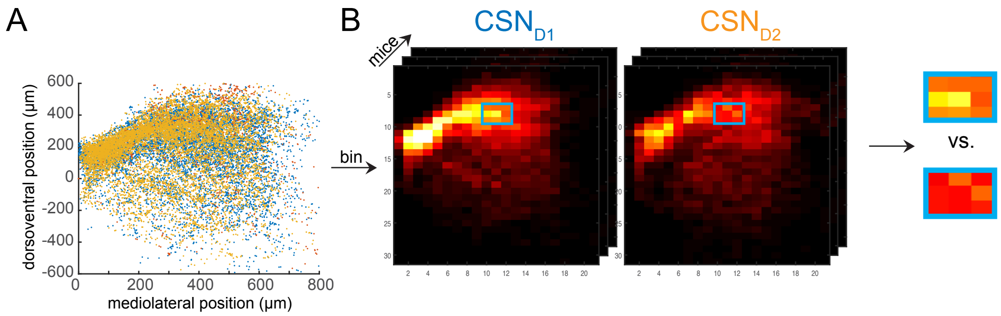
A
B
dorsoventral position (µm)
mediolateral position (µm)
CSN$_{D1}$
CSN$_{D2}$
mice
bin
vs.

Figure S5

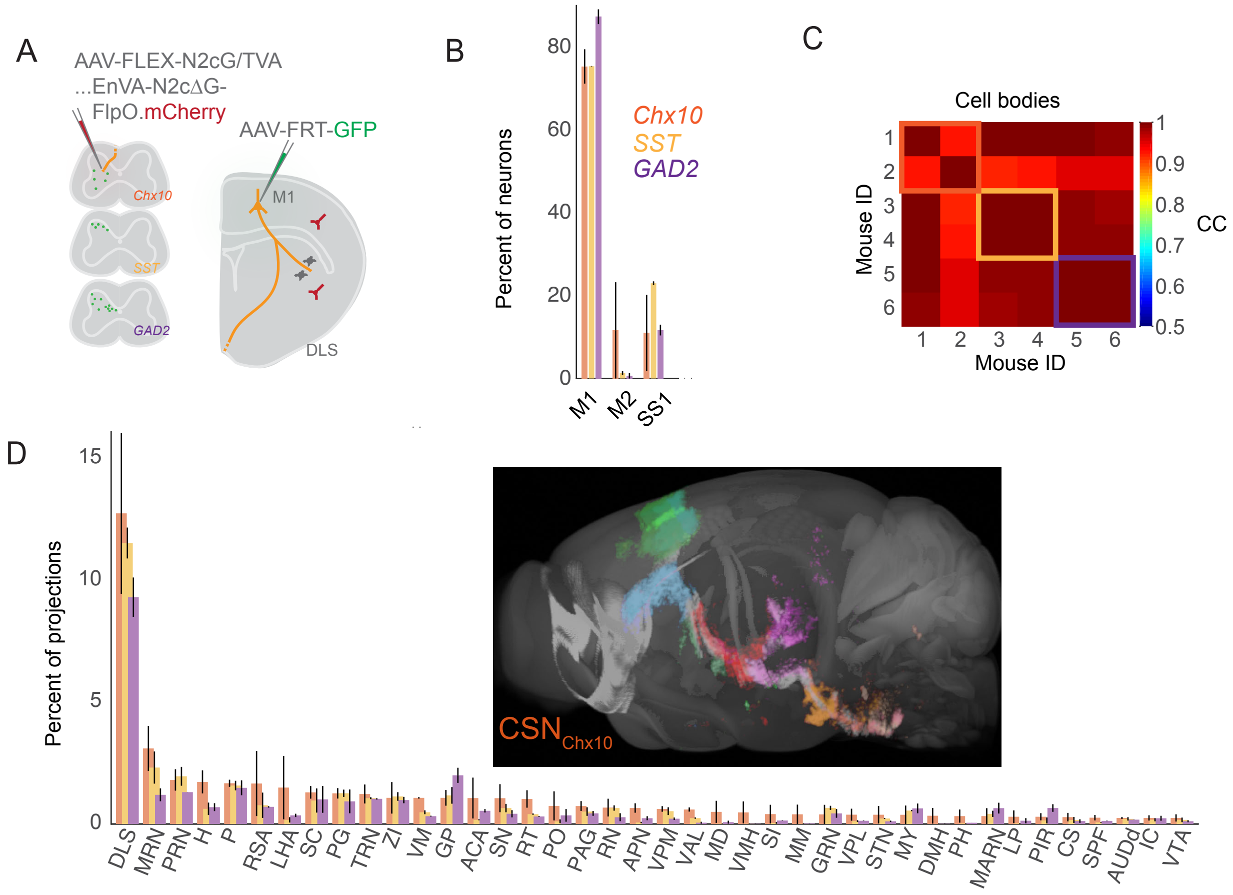
A
AAV-FLEX-N2cG/TVA
...EnVA-N2cΔG-
FlpO.mCherry
AAV-FRT-GFP
Chx10
SST
GAD2
M1
DLS
B
Percent of neurons
0
20
40
60
80
M1
M2
SS1
Chx10
SST
GAD2
C
Cell bodies
Mouse ID
1
2
3
4
5
6
CC
1
0.9
0.8
0.7
0.6
0.5
D
Percent of projections
0
5
10
15
CSN<sub>Chx10</sub>
DLS MRN PRN H P RSA LHA SC PG TRN ZI VM GP ACA SN RT PO PAG RN APN VPM VAL MD VMH SI MM GRN VPL STN MY DMH PH MARN LP PIR CS SPF AUDd IC VTA

Figure S6

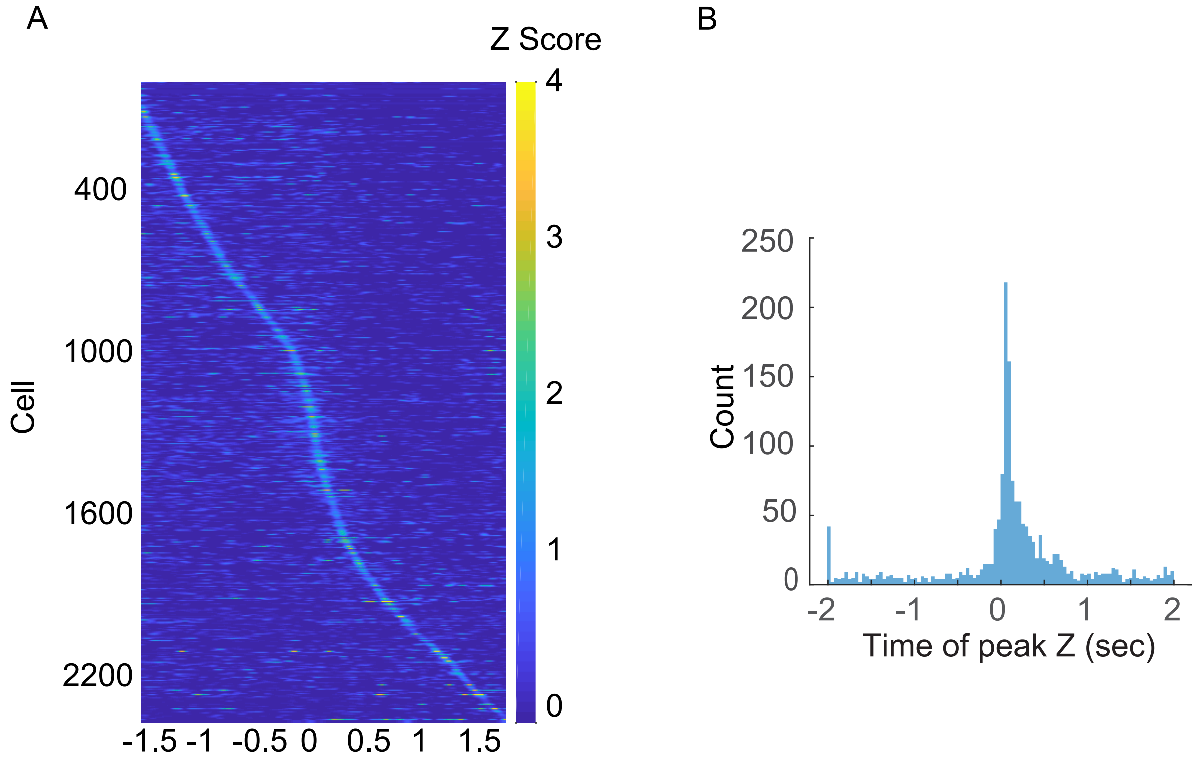
A
Z Score
4
3
2
1
0
Cell
400
1000
1600
2200
-1.5 -1 -0.5 0 0.5 1 1.5
B
250
200
150
100
50
0
Count
-2 -1 0 1 2
Time of peak Z (sec)

Figure S7

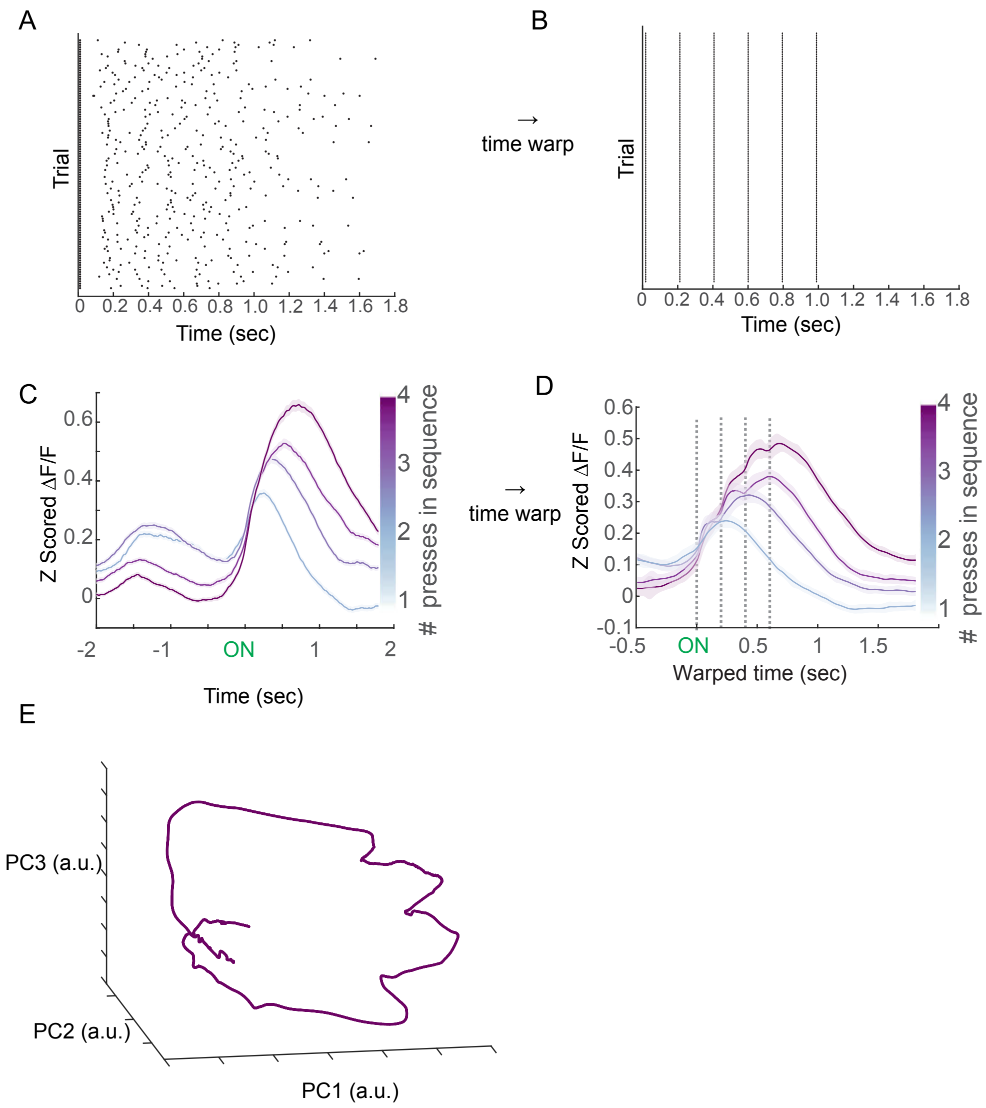
A
Trial
0 0.2 0.4 0.6 0.8 1.0 1.2 1.4 1.6 1.8
Time (sec)
→
time warp
B
Trial
0 0.2 0.4 0.6 0.8 1.0 1.2 1.4 1.6 1.8
Time (sec)
C
Z Scored ΔF/F
0.6
0.4
0.2
0
-2 -1 ON 1 2
Time (sec)
4 3 2 1
### presses in sequence
→
time warp
D
Z Scored ΔF/F
0.6 0.5 0.4 0.3 0.2 0.1 0 -0.1
-0.5 ON 0.5 1 1.5
Warped time (sec)
4 3 2 1
### presses in sequence
E
PC3 (a.u.)
PC2 (a.u.)
PC1 (a.u.)

Figure S8

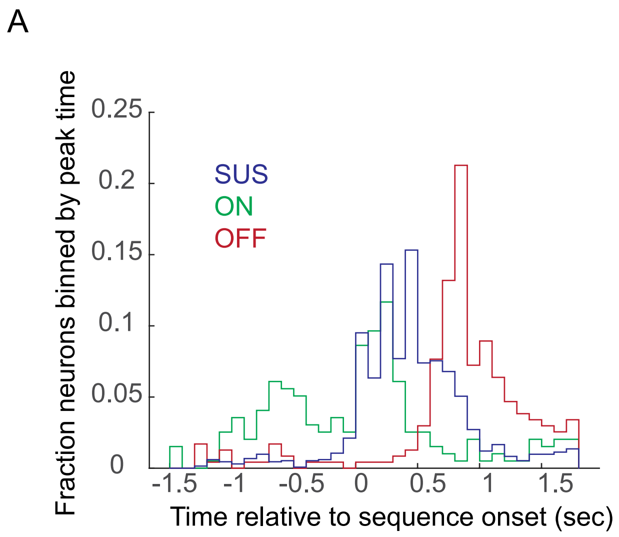
A
SUS
ON
OFF
Fraction neurons binned by peak time
0.25
0.2
0.15
0.1
0.05
0
-1.5
-1
-0.5
0
0.5
1
1.5
Time relative to sequence onset (sec)

Figure S9

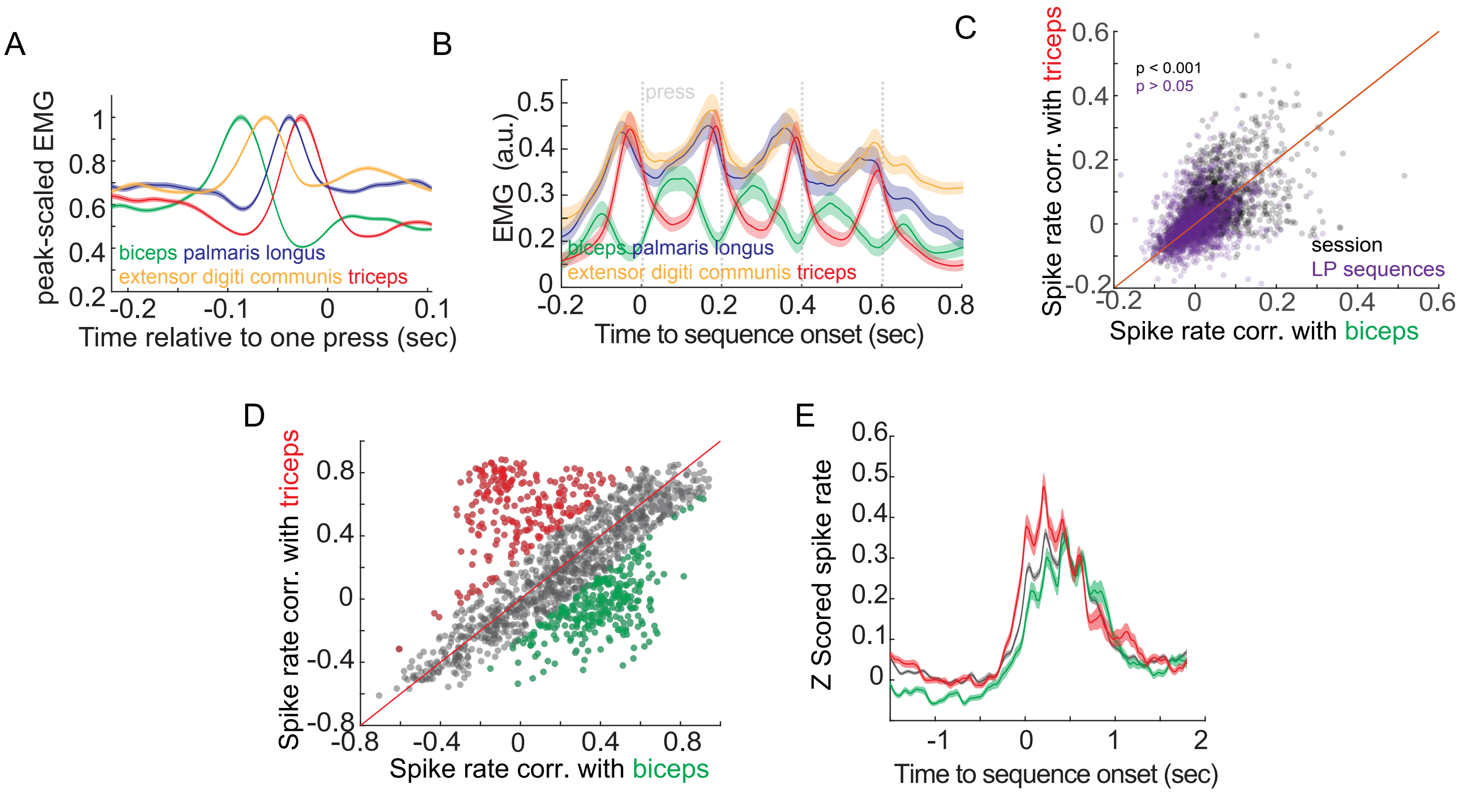
A
peak-scaled EMG
Time relative to one press (sec)
biceps palmaris longus
extensor digiti communis triceps
B
EMG (a.u.)
press
Time to sequence onset (sec)
biceps palmaris longus
extensor digiti communis triceps
C
Spike rate corr. with triceps
Spike rate corr. with biceps
p < 0.001
p > 0.05
session
LP sequences
D
Spike rate corr. with triceps
Spike rate corr. with biceps
E
Z Scored spike rate
Time to sequence onset (sec)

Figure S10

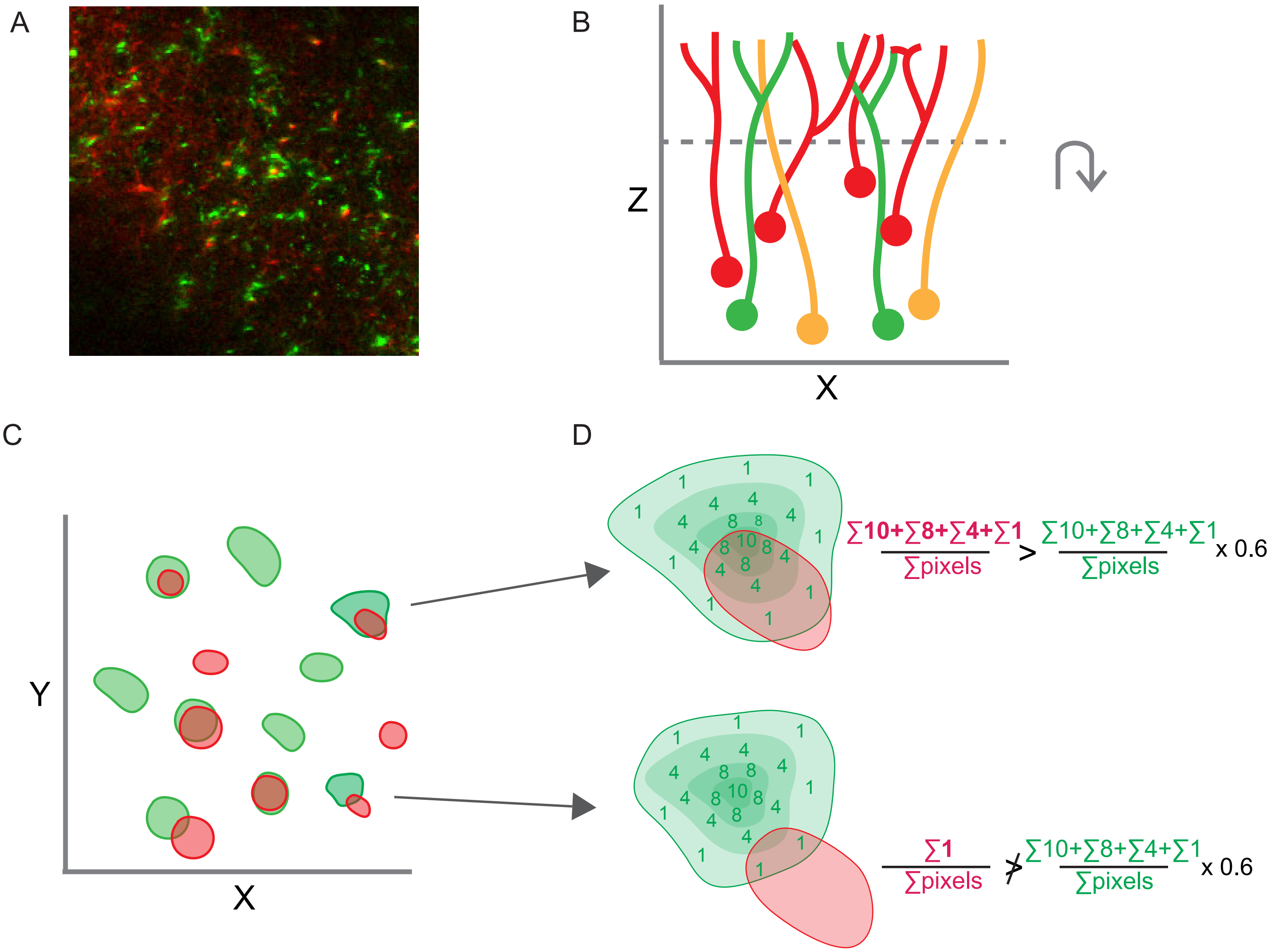

A
B
Z
X
C
Y
X
D
∑10+∑8+∑4+∑1
∑pixels
>
∑10+∑8+∑4+∑1
∑pixels
x 0.6
∑1
∑pixels
≯
∑10+∑8+∑4+∑1
∑pixels
x 0.6


Figure S11
